# The Evolution of Extreme Genetic Variability in a Parasite-Resistance Complex in a Planktonic Crustacean

**DOI:** 10.1101/2024.08.09.607325

**Authors:** Suha Naser-Khdour, Fabian Scheuber, Peter D. Fields, Dieter Ebert

## Abstract

Genomic regions that play a role in parasite defense are often found to be highly variable, with the MHC serving as an iconic example. Single nucleotide polymorphisms may represent only a small portion of this variability, with Indel polymorphisms and copy number variation further contributing. In extreme cases, haplotypes may no longer be recognized as homologs. Understanding the evolution of such highly divergent regions is challenging because the most extreme variation is not visible using reference-assisted genomic approaches. Here we analyze the case of the Pasteuria Resistance Complex (PRC) in the crustacean *Daphnia magna*, a defense complex in the host against the common and virulent bacterium *Pasteuria ramosa*. Two haplotypes of this region have been previously described, with parts of it being non-homologous, and the region has been shown to be under balancing selection. Using pan-genome analysis and tree reconciliation methods to explore the evolution of the PRC and its characteristics within and between species of *Daphnia* and other *Cladoceran* species, our analysis revealed a remarkable diversity in this region even among host species, with many non-homologous hyper-divergent-haplotypes. The PRC is characterized by extensive duplication and losses of Fucosyltransferase (FuT) and Galactosyltransferase (GalT) genes that are believed to play a role in parasite defense. The PRC region can be traced back to common ancestors over 250 million years. The unique combination of an ancient resistance complex and a dynamic, hyper-divergent genomic environment presents a fascinating opportunity to investigate the role of such regions in the evolution and long-term maintenance of resistance polymorphisms. Our findings offer valuable insights into the evolutionary forces shaping disease resistance and adaptation, not only in the genus *Daphnia,* but potentially across the entire *Cladocera* class.

**Significance:** Understanding how organisms adapt to their environment requires insights into the evolution of genetic defenses against their parasites. While the Major Histocompatibility Complex (MHC) is a well-known example of a highly variable immune-related gene region, much remains unknown about the evolution of other such regions. Our study investigates the Pasteuria Resistance Complex (PRC) in water fleas, a genomic region crucial for defense against a parasitic bacterium. We discovered that the PRC is exceptionally diverse, with a history spanning hundreds of millions of years. This research provides new insights into the mechanisms underlying the maintenance of genetic diversity in the face of persistent parasite pressure. Our findings contribute to a broader understanding of how organisms evolve robust defenses against infectious diseases.

## Introduction

For decades, scientists have been grappling with the question of how highly polymorphic genomic regions are maintained in populations. Because their genetic diversity is much higher than the genomic average of a population, these regions stand out. The best known example and the main model system for studying diverse genomic regions and their evolution is the major histocompatibility complex (MHC) of jawed vertebrates (Piertney and Oliver 2006; Radwan, et al. 2020). The MHC is composed of tightly linked loci, that collectively govern complex traits with inheritance patterns equivalent to alternative alleles at a single locus (Thompson and Jiggins 2014). The origin of such genomic regions is intricately linked to traits that require coordination among multiple genes, so natural selection plays a crucial role in their evolution, favoring combinations of alleles across genes that confer advantageous traits. Alternative haplotypes may be maintained by balancing selection (Schwander, et al. 2014; Thompson and Jiggins 2014; Gutierrez-Valencia, et al. 2021; Berdan, et al. 2022), but over time, they may undergo further evolutionary changes as new mutations become integrated, expanding their functional scope and driving their genetic divergence.

The maintenance of highly polymorphic regions relies heavily on the reduction or even suppression of recombination between the linked loci (Charlesworth, et al. 2005). Reduced recombination ensures that specific combinations of alleles are maintained, preventing less fit recombinant haplotypes from forming (Yeaman 2013). Suppression of homologous recombination is often associated with inversions, although mechanisms such as large insertions, deletions (Gutierrez-Valencia, et al. 2021), or duplications (Kulski, et al. 2022) can also be involved.

One explanation for high variability is balancing selection. In the context of pathogen-resistance loci, balancing selection can prevent the fixation of alleles, resulting in the persistence of multiple variants with different resistance phenotypes against diverse parasite strains (Haldane 1992; Paterson, et al. 1998; Thrall, et al. 2012; Chiarella, et al. 2023). This scenario can lead to negative frequency-dependent selection (NFDS), in which rare alleles have an advantage, increasing their frequency at the population level, and thus promoting the maintenance of diverse haplotypes (Tellier, et al. 2014; Ebert and Fields 2020). Interestingly, a recent discovery in the *Caenorhabditis elegans* nematode identifies the presence of punctuated regions of extreme divergence, called hyper-divergent-haplotypes, potentially maintained by long-term balancing selection (Lee, et al. 2021). A region of extreme divergence has earlier been reported in the crustacean *Daphnia*, linked to pathogen resistance (Bento, et al. 2017) and shown to be under balancing selection (Bourgeois et al. 2021). Here we further investigate this region in *Daphnia*, aiming to better understand the evolution and the characterising features of such hyper-divergent-haplotypes.

The Pasteuria Resistance (PR) locus of the planktonic crustacean *Daphnia magna* presents a fascinating departure from the classical definition of supergenes. Initially this locus was described as a supergene with three tightly linked resistance loci that had a significantly reduced recombination rate (Bento, et al. 2017) and played an important role in defense against the virulent parasite *Pasteuria ramosa* (Ameline, et al. 2021). Unlike most supergenes, however, the PR locus contains no inversions to suppress recombination; rather, its low recombination rate may be related to the very high divergence of haplotypes (Bento, et al. 2017). We refer here to the PR locus and two other more recently discovered resistance loci, the nearby F (Fredericksen, et al. 2023) and G-loci (Eglantine, in prep.), as the Pasteuria Resistance Complex (PRC). The remarkable polymorphisms observed for resistance phenotypes at this complex raise intriguing questions about the evolutionary forces shaping this unusual pattern of variation.

In this study we will first explore the extent of variation in the PRC by investigating its variation in sequence composition, gene content, and overall architecture both within populations of *Daphnia magna* and across related species. Furthermore, we explore the origin of the PRC by using tree reconciliation (Page 1994) to compare species phylogeny with gene phylogeny. To characterize the gene content of the PRC, we mainly focus on fucosyltransferase genes, which are present in large numbers in the PRC and have been suggested as strong candidates for explaining polymorphism in the resistance loci within the PRC (Bento, et al. 2017; Fredericksen, et al. 2023) (& Eglantine’s paper). In addition, we investigate the potential influence of the HDR on the evolution and structure of the PRC and how this affects the long-term role of this complex in *Daphnia’s* defense mechanisms. Through this comprehensive investigation, we hope not only to illuminate the evolution of the *Daphnia magna* PRC, but also contribute to a broader understanding of resistance loci, particularly within the context of hyper-divergent regions.

## Results

### Identifying the PRC in *D. magna*

Exploration of the PRC across many Daphnia genotypes confirmed previous observations about the substantial size and content variation seen between two *Daphnia magna* haplotypes (Bento et al., 2017).

For this work, we began by using the PRC described by Bento et al. (2017) and Fredericksen et al. (2023) to locate the PRC in an additional 13 long-read-based genomes from *D. magna* clones using BLASTN, and then subsequently constructed a pan-genome graph with PanGraph (Noll et al. 2023) for this region. Our analysis revealed multiple conserved sequences at the start of the PRC and in the region containing the F-locus (Fig. 1A; Supplementary Fig. 1). However, because each clone’s PRC exhibited a unique non-homologous haplotype of varying lengths (Fig. 2; Supplementary Fig. 1), the proportion of regions that could be aligned using all 14 *D. magna* clones was rather low, ranging from 27% to 55% with a mean of 49% (Supplementary Fig. 1). In addition, the pairwise pangenome graphs of all *D. magna* clones (total of 91 pan-genome graphs), showed that the proportion of alignable regions ranged between 31% and 93%, with a mean of 55% (Supplementary Table 1), indicating that the mean proportion of homologous regions is similar whether it is a pair-wise comparison or an all-clones comparison.

**Fig. 1.**
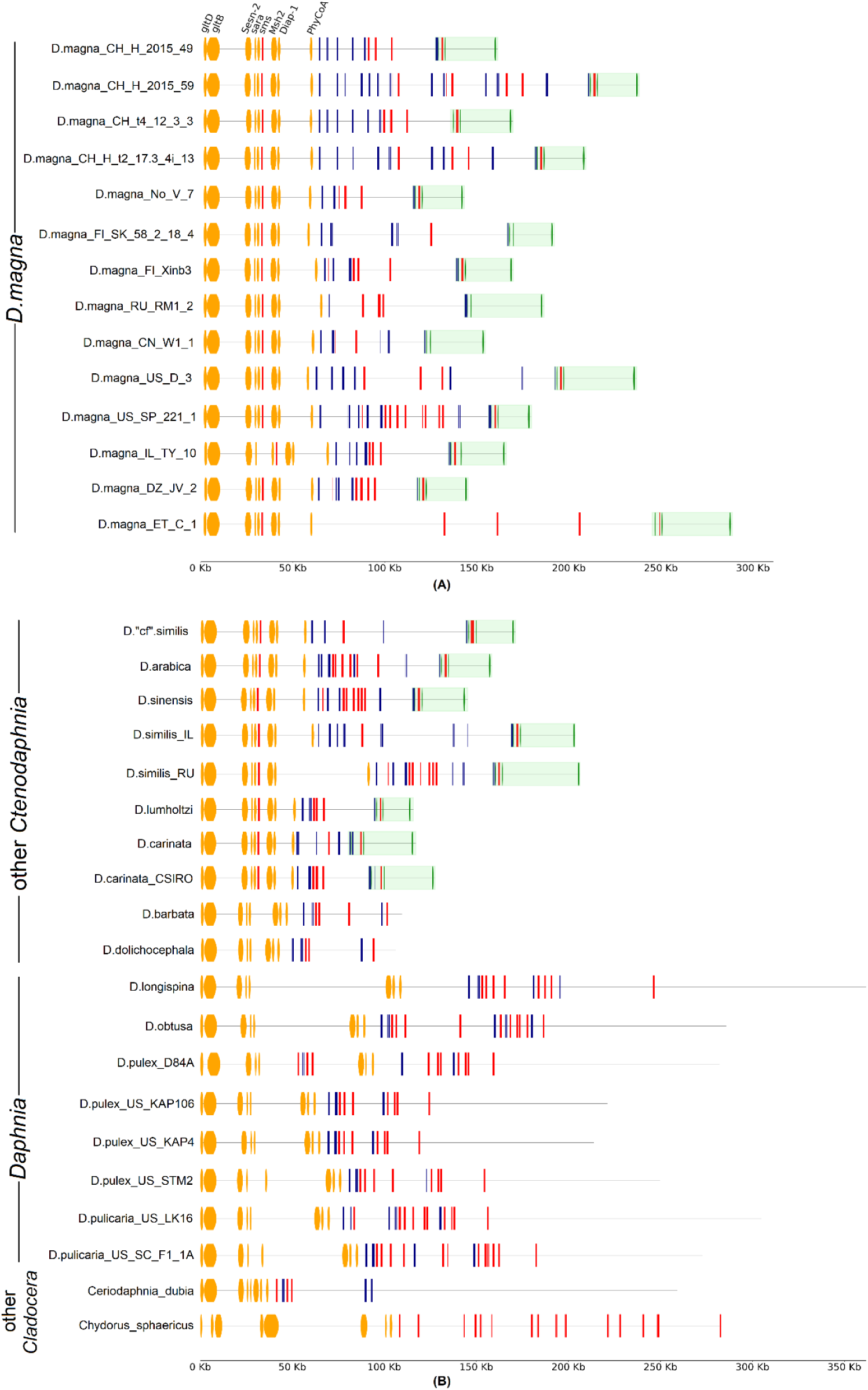
The PRC is characterized by a conserved region across diverse taxa, followed by a highly polymorphic FuT island (genes in red and blue). A) PRC alignment of 14 clones of *D. magna*. B) PRC alignment of other Cladoceran species. Color legend: Orange: conserved region across all taxa, Green: conserved region across *Ctenodaphnia* (ex. *D. barbata* and *D*.

**Fig. 2.**
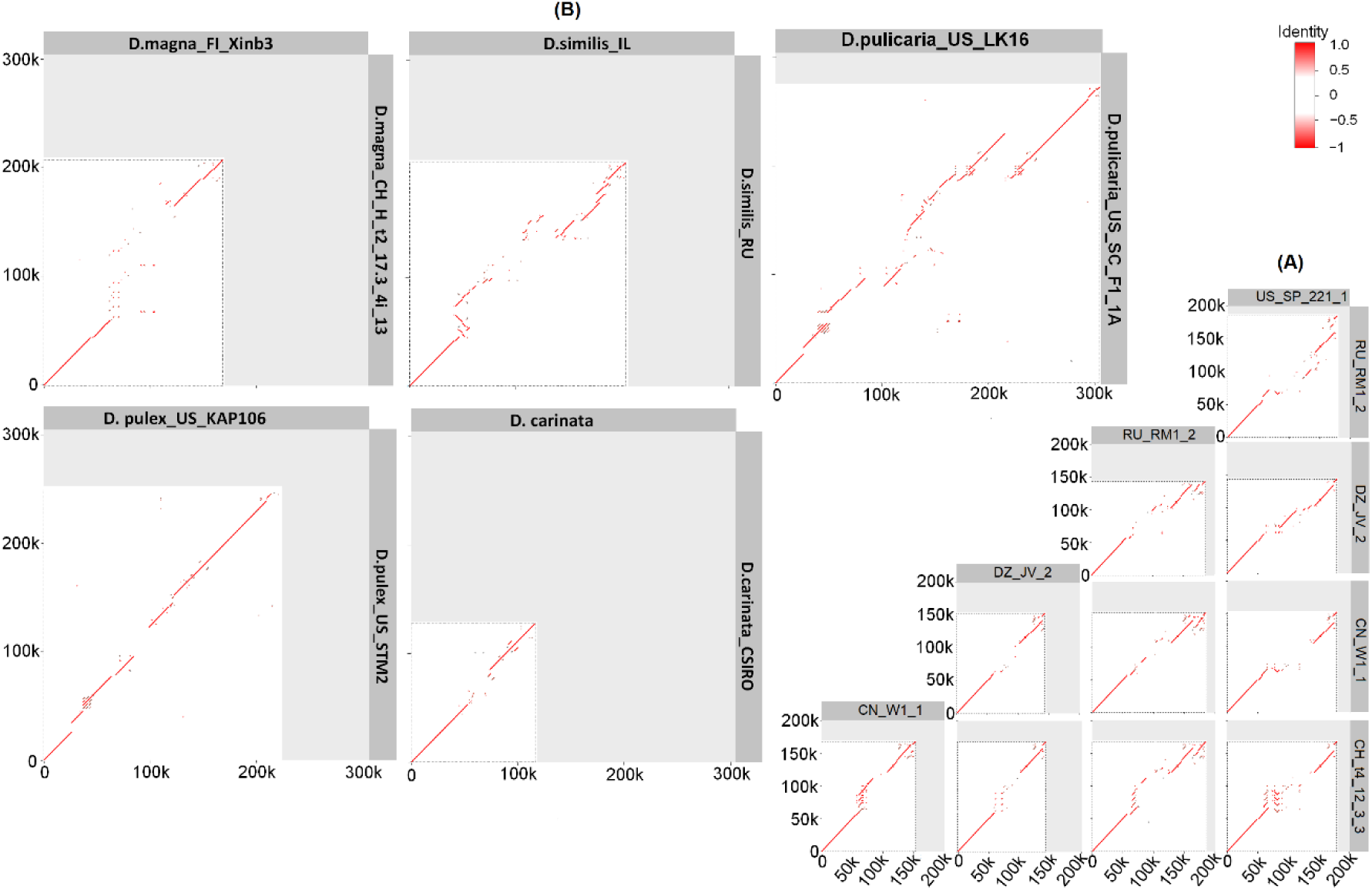
Pairwise dotplots of the PRC and flanking regions reveal the presence of non-homologous regions within several *Daphnia* species. A) Examples of dotplots between pair of *D. magna* clones, and B) between pairs of clones from 5 different species.

We characterized the gene content in both conserved and non-homologous regions of all genomes independently using ORF Finder and BLASTX. The non-homologous region consisted primarily of multiple duplicates of fucosyltransferase genes (FuT) and, often, galactosyltransferase (GalT) genes, forming a distinct “FuT island” (Fig. 3A) that differed among genotypes. In contrast, the conserved region at the beginning of the PRC harboured genes annotated as important for cell growth (Fig. 3A ; Fig. 1A, orange regions), including Glutamate synthase subunits (gltD, gltB), Sestrine-2 (Sesn-2), Sara protein (sara), Spermine synthase (sms), DNA mismatch repair protein (Msh2), Death-associated inhibitor of apoptosis 1 (Diap1), and Phytanoyl Coenzyme A (PhyCoA) (de Wind, et al. 1995; Wang, et al. 1999; Kershaw, et al. 2001; Jönsson and Lowther 2007; Pegg and Michael 2010; Liu and Jin 2017; Loubery, et al. 2017; Lu, et al. 2019; Wang, et al. 2023). Notably, the conserved F-locus regions (Fig. 1A, green regions) mainly consisted of non-coding sequences and some uncharacterized genes that belong to a family of Cladoceran-specific genes (for the list of genes from two haplotypes see Bento, et al. 2017 and Fredericksen, et al. 2023).

**Fig. 3.**
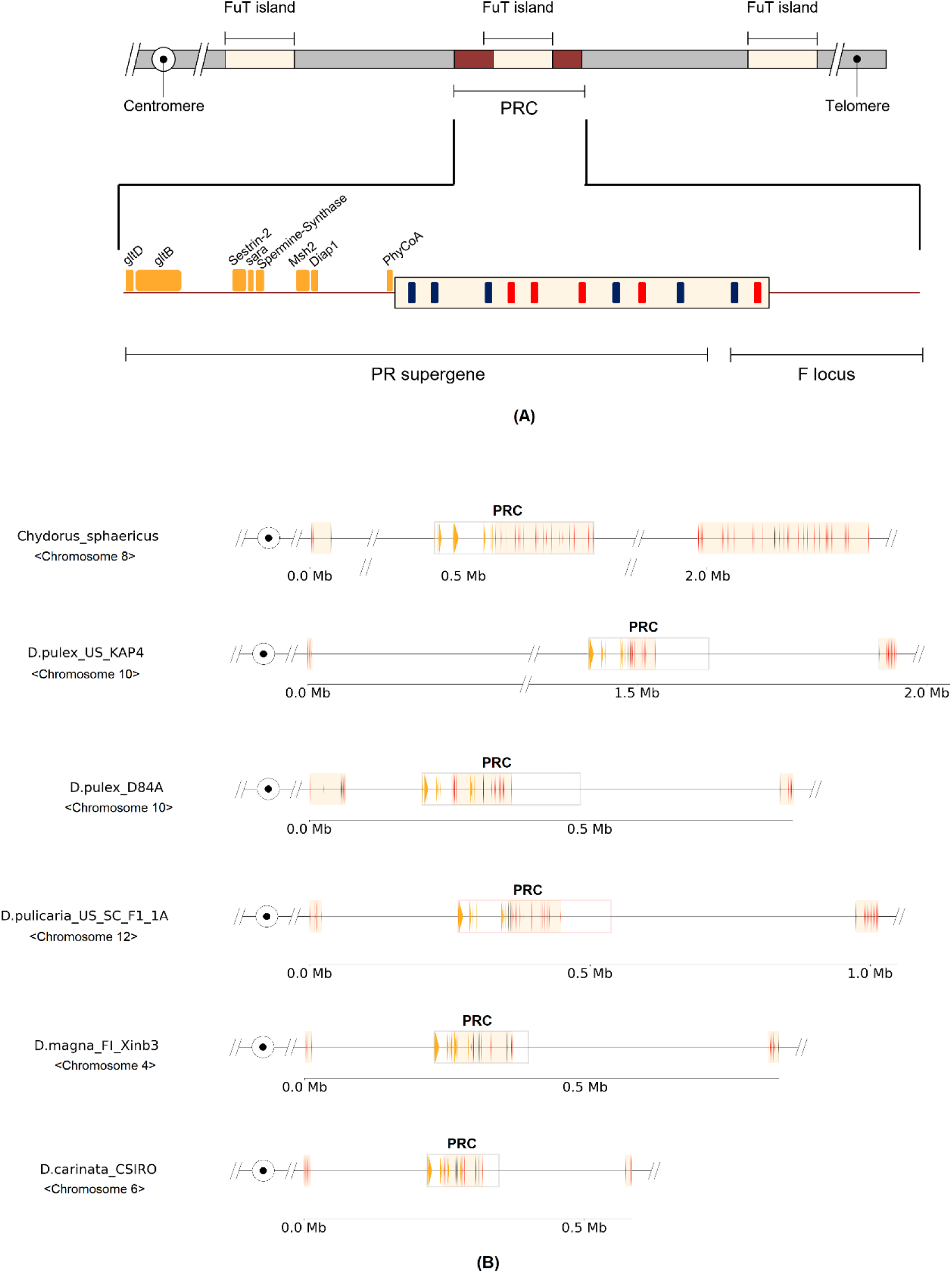
A) Schematic representation of the PRC. Eight conserved genes were used to identify the complex (orange) and the FuT island with multiple copies of FucT (red) and GalT (blue). B) PRC alignment of six species for which we have a chromosome-level assembly and that include the two FuT islands surrounding the PRC.

### Analysis of the PRC species in other species

We also searched for the PRC in other *Daphnia* and related species where high quality genomes were available; however, the high variability of the PRC posed a challenge in assembly, particularly due to the complexity of the FuT island. Therefore, we incorporated into our analysis only assemblies generated from long reads that contained the PRC within a single contig. From 45 high quality assemblies examined (Supplementary Fig. 2; Supplementary Table 2), 20 non-*D. magna* assemblies that contained the entire PRC in one contig were identified.

These genome assemblies all shared the same structure of the PRC seen in *D. magna* (Fig. 1B), including the genome of *Chydorus sphaericus*. Thus, we can trace the origin of the PRC to a common ancestor of *Daphnia* and *Chydorus sphaericus* more than 250 million years ago (van Damme et al 2022). Extending our search to crustaceans outside the *Cladocera* (*Leptestheria dahalacensis*, *Eulimnadia texana*, *Lepidurus packardi*, *Capitulum mitella*, *Penaeus monodon*, and *Tigriopus japonicus*) that shared a common ancestor with *Daphnia* even longer ago, we found no discernible traces of the PRC.

In general, the overall structure of the PRC appears highly conserved with only a few rearrangements observed among the eight conserved genes. The conserved region of the F locus could not be detected outside the *Ctenodaphnia* clade, and within this sub-genus, the F locus was identified in all *Ctenodaphnia* species except in the branch leading to *D. barbata* and *D. dolichocephala* (Fig. 1B), a more ancestral branch of the sub-genus. Thus, the F locus emerged as part of the PRC after the divergence of the *D. barbata* - *D. dolichocephala* clade from the other *Ctenodaphnia* species included here.

### FuT islands in the PRC region

An interesting feature of the PRC is the FuT island, which is characterized by extensive duplication of FuT genes and often GalT genes, with variability in gene number and position even among genotypes within the same population (Fig. 1, the 4 *D. magna* clones starting with CH are from the same population). We define the FuT island as a genomic region where at least three copies of FuT/GalT and at least one copy of FuT reside within a distance of 50 kb from each other. FuT and GalT are suspected to play a major role in defending against *P. ramosa* attachment in *D. magna* (Bento, et al. 2017; Fredericksen, et al. 2023). The FuT islands’ outstanding polymorphism may correspond with the PRC’s high diversity for resistance polymorphism. Moreover, the polymorphism seen in *D. magna*, extends to other *Daphnia* species where over one high quality genome was available, such as the closely related *D. similis* and *D. carinata* and the more distantly related *D. pulex* (Fig. 2B), suggesting that this polymorphism dates back to at least 145 Mya (Cornetti, et al. 2019).

The FuT islands, while associated with the PRC, are not exclusive to it. Previous research on the North American *D. pulex* genome has revealed a significant expansion of the FuT gene family compared to other non-Crustacean genomes (Colbourne, et al. 2011). Our findings indicate widespread presence and strong clustering of FuT copies in Crustacean; in the investigated species, over 75% of FuT’s were located on three chromosomes, one of which contained the PRC (Fig. 3B; Supplementary Fig. 3). Apart from the FuT island within the PRC, two similar islands situated approximately 300 kbp away (median = 257,409 bp, mean = 377,865 bp) flank the complex. These islands have on average less FuT copies than the PRC (Supplementary Table 2). There are no FuT genes present between these flanking islands and the PRC (Fig. 3A). These results suggest a distinctive genomic arrangement characterized by clustering of FuT copies within specific chromosomal regions, including the PRC and adjacent to it (Fig. 3B; Supplementary Fig. 4).

### The highly polymorphic PRC has a single ancestral origin

To explore the evolutionary origin of the PRC, we used 45 available *Crustaceans* genomes, plus 14 new draft genomes from 13 different non-*D.magna*, *Daphniidae* species (Supplementary Table 3, Supplementary Fig. 2). Using the non-reversible model (Minh, et al. 2020), we inferred the rooted species tree from 90 BUSCO genes with 89.5% bootstrap support (Naser-Khdour, et al. 2021) for the root on the divergence node between the *Multicrustacea* and the *Branchiopoda* clades (Fig. 4A; Supplementary Fig. 5A). Besides some nodes within the *D.magna* clade, all nodes in the species tree have high bootstrap support values (Minh, et al. 2013).

**Fig. 4.**
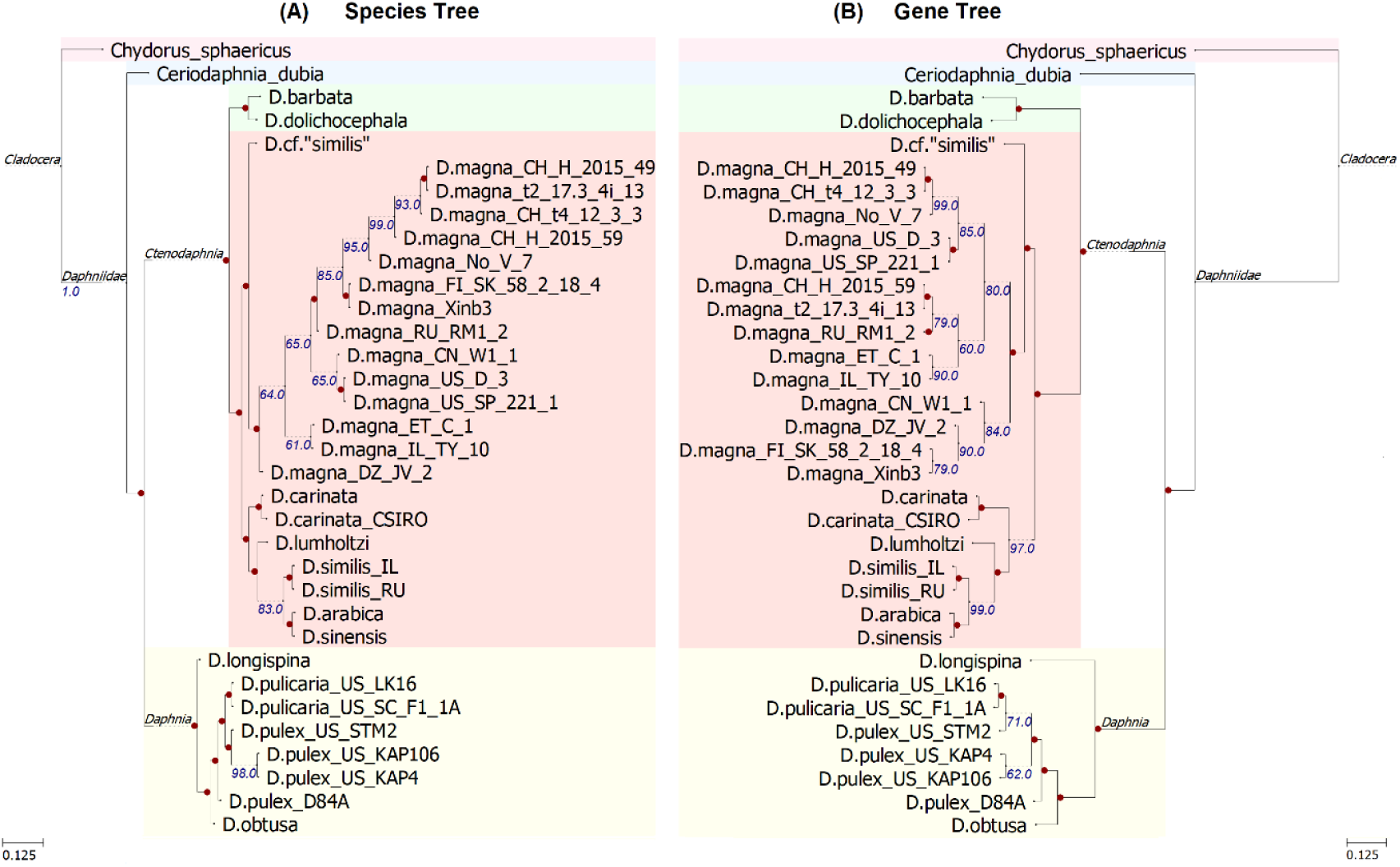
ML phylogeny based on A) arthropod BUSCO genes and B) the eight conserved genes from the PRC.

Using the eight conserved genes in the PRC (Fig. 1), we inferred the ML gene tree based on the concatenated alignment of those genes and compared it to the ML species tree (Fig. 5). In addition, we inferred the coalescent-based gene tree from the individual gene trees. Our results show that the ML gene tree and species tree are congruent and consistent with the coalescent-based phylogeny (Supplementary Fig. 5B).

**Fig. 5.**
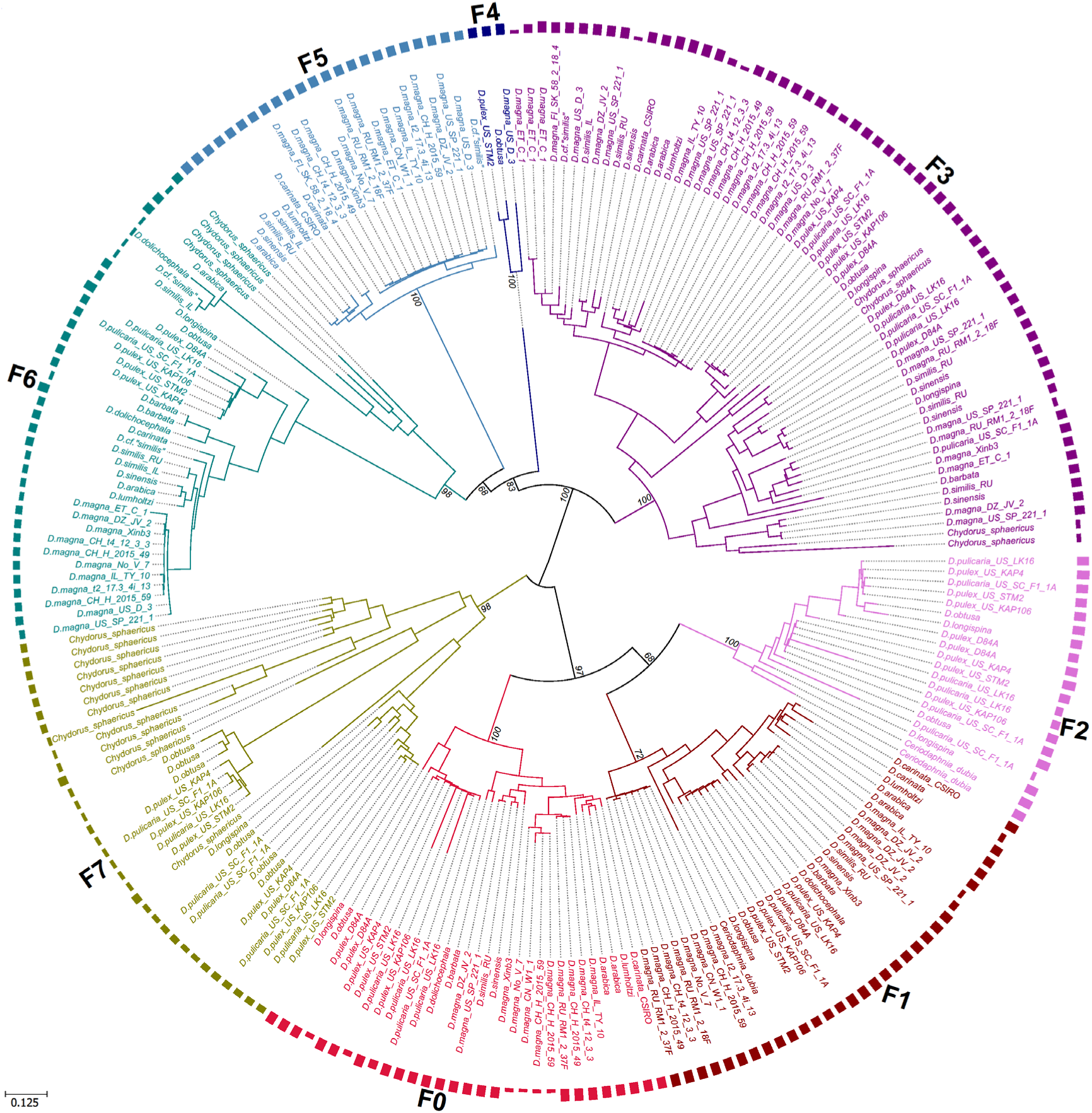
ML FuT tree, based on FuT copies from all identified PRC in 34 taxa. The bars in the outer circle correspond to the length of each FuT copy.

### The FuT gene tree shows a different evolutionary history from the PRC gene tree

All FuT copies observed in the PRC were one-exon genes. Using all FuT genes from the PRC of all species, we generated the gene tree of all 35 taxa with PRC using IQTREE2 (Minh, et al. 2020).

We found eight distinctive clades of FuT, arbitrarily denoted here as F0, F1, …, F7. To qualify as a clade, FuT sequences needed to be consistently monophyletic in both the amino acid and nucleotide trees, and the branch leading to the clade had to be greater than 0.3. This approach revealed some clades to be consistent with the species tree, but others not (Fig. 5; Supplementary Fig. 6). For example, F2 and F7 clades include no *Ctenodaphnia* subgenus species, yet all species from the subgenus *Daphnia* have FuT duplicates that belongs to those clades (Fig. 5; Supplementary Fig. 6). Overall, the gene tree of FuT duplicates shows a different evolutionary history than the conserved region of the PRC and the species tree, mainly due to duplications and loses of FuT copies.

We found that three FuT clades are shared with *Chydorus sphaericus* (Fig. 5; Supplementary Fig. 5, clades - F3,F6 and F7), and two clades shared with *Ceriodaphnia dubia* (Fig. 5; Supplementary Fig. 6 clades - F1 and F2), a *Daphniidae* species that diverged from the *Daphnia* genus around 153 Mya (Van Damme, et al. 2022). The other three FuT clades were either exclusive to the *Daphnia* genus (Fig. 5; Supplementary Fig. 6, clade F0), to the *Ctenodaphnia* subgenus (Fig. 5; Supplementary Fig. 6, clade F5), or exclusive only to a few clones from different species (Fig. 5; Supplementary Fig. 6, clade F4).

To understand the evolutionary history of the FuT gene in the PRC, we reconciled the FuT gene tree with the ML species tree and found many gene duplications (119) throughout the evolutionary tree, but even more loses of FuT copies (476 loses), especially in the *Ctenodaphnia* clade (Fig. 6). Furthermore, when we redid the reconciliation of the FuT gene tree with the ML species tree, treating each of the 8 FuT clades as an independent gene family, the results were comparable, albeit with slightly fewer duplications and loses in the analysis of the subdivided FuT gene set (Supplementary Table 4). This analysis of FuT evolution sheds light on the complex dynamics of this gene family, highlighting both duplicative events and losses, and helping to explain the high variability of the PRC among genotypes from within and between species.

**Fig. 6.**
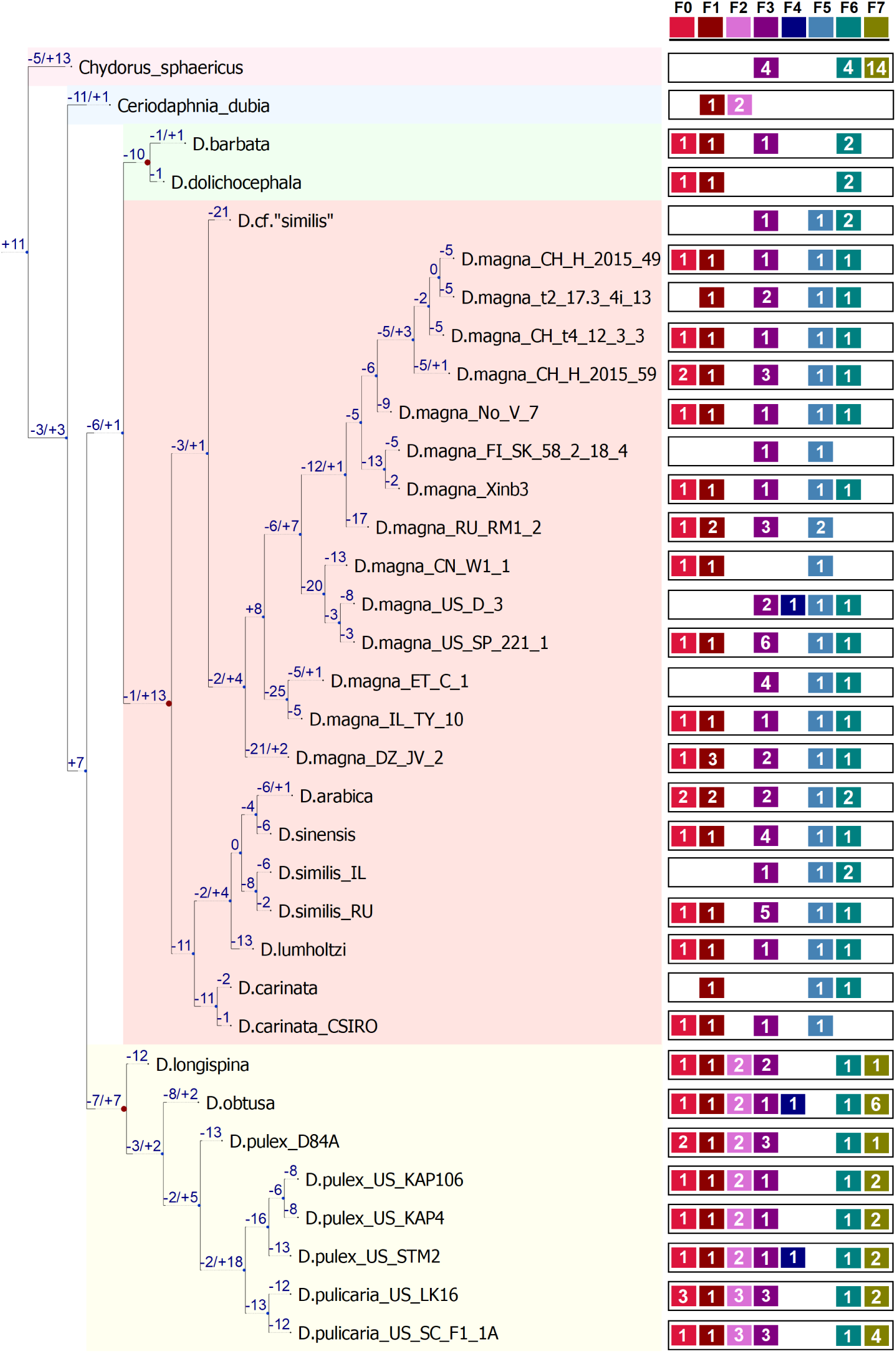
A phylogenetic reconciliation between species tree and FuT gene tree with the number of FuT duplications (+) and loses (-) on each branch

## Discussion

With the increased use of long-read next generation sequencing, more and more structural variation is being discovered in genomes of diverse organisms, and extreme cases of hyper-divergent haplotypes are being noted (Lee, et al. 2021). Yet, very little is known about the structure and evolution of such regions. Based on previous findings about the genetic and phenotypic diversity of the *D. magna* resistance gene complex under balancing selection, our deeper investigation into its structure and evolution found enormous structural genetic diversity, both within populations, species and across related species. The hyper-divergent-haplotypes of this resistance complex exhibit extensive non-homologous regions reaching up to 100 kb, and, unlike classical supergenes, where typically only few haplotypes are segregating, each haplotype appears unique. This remarkable diversity can be attributed, at least in part, to the absence of recombination within this region. The Messelson Effect (Mark Welch and Meselson 2000) predicts that asexually reproducing allelic copies at a given locus, like the PRC, accumulate mutations independently, providing a framework for understanding the evolution of the observed extreme divergence between haplotypes..

Balancing selection is a likely driver in maintaining the exceptional polymorphism of the PRC. This mode of selection favors the persistence of multiple haplotypes by preventing rare haplotypes to be lost. At the same time, it fosters the accumulation of neutral and deleterious mutations, i.e. “Sheltered load” (Uyenoyama 1997, 2005). However, it is unlikely the driver of the PRC’s initial evolution. The mechanisms underlying the PRC’s origin remain unclear, with no evidence suggesting that arose from mechanisms frequently discussed, like inversions or horizontal gene transfer. A speculative hypothesis proposes that the PRC originated from two closely linked polymorphic loci involved in pathogen defense. Certain allele combinations at these loci may have conferred a selective advantage, while others were detrimental. To preserve advantageous combinations, suppression of recombination between these loci became beneficial, leading to the formation of the ancestral PRC. Over time, this initial structure expanded through the accumulation of mutations, including gene duplications, insertions, and deletions, resulting in the observed hyper-divergent haplotypes. Additionally, rare recombination events, although suppressed, might have further shuffled existing variation, contributing to the current repertoire of haplotypes.

We were able to trace the evolution of this PRC complex back to a common ancestor over 250 million years ago. Characteristically, the highly variable part of this resistance complex contains vast structural variation among members of the fucosyltransferase gene family, a group of genes suspected to play a role in the resistance mechanism of this complex. Gains and losses of FuT gene family members are apparent throughout the *Daphnia* clade, explaining part of this region’s high variability. Our investigation unveils novel insights into the evolution and maintenance of the *Daphnia magna* PRC, particularly its remarkable antiquity and unique spatial context within a hyper-divergent region.

Our findings support the presence of the PRC in diverse *D. magna* populations, largely extending earlier observations by Bento et al. (2017). Consistent with their work, we observed substantial variation in PRC size and content across *Daphnia* genotypes within and between species, highlighting its dynamic nature and potentially reflecting the coevolution between *D. magna* and its pathogen *P.ramosa*, for which this system is well known (Decaestecker, et al. 2007; Ameline, et al. 2021; Bourgeois, et al. 2021). Moreover, our results show that the presence of the PRC extends way beyond *D. magna*, possibly having emergered in the *Anomopoda* order or even the *Cladocera* superorder, which date it back a remarkable 250-400 million years ago.

The comparative analysis revealed several conserved sequences in the PRC at the beginning of the region: these likely harbour essential roles for the animals’ overall function unrelated to the PRC’s known function of resistance. Conversely, the presence of extensive non-homologous regions across different clones and taxa suggests rapidly evolving sequences that may contribute to pathogen resistance specificity. This discovery aligns with theoretical predictions that high polymorphism offers long-term adaptive advantages (Dobzhansky and Pavlovsky 1960). The persistence of hyper-divergent haplotypes in the PRC through numerous speciation events provides a striking example of a polymorphism that is likely maintained by host-parasite co-evolution (Cornetti, et al. 2024).

The observed variation within the PRC raises intriguing questions about the underlying mechanisms driving its evolution. The incongruence of species and FuT gene trees suggest that negative-frequency-dependent selection, where rarer resistance alleles gain an advantage as the common ones become targeted by pathogens, might help maintain this diversity (Ebert and Fields 2020). In addition, the location of this complex within a hyper-divergent region presents a unique opportunity to explore the potential relationship between such regions and the evolution of pathogen-resistance loci. Even though hyper-divergent regions have been reported in other species, e.g. nematodes (Lee, et al. 2021) and on other *D. magna* chromosomes (Dexter et al. 2024. Manuscript in preparation), the remarkable diversity observed in the PRC itself, including the distinct “FuT island”, suggests the potential for hyper-divergent regions to harbor functionally important genetic variations.

Our discovery of multiple FuT islands within the hyper-divergent region (HDR) on chromosome 4 strengthens the link between glycosylation and parasite resistance observed in the *D. magna* F-locus study (Fredericksen et al. 2023). These FuT islands potentially encode a diverse repertoire of α-(1,3)-fucosyltransferases, Glycoprotein-N-acetylgalactosamine-3-β-galactosyltransferases, α-(1,4)-N-acetylglucosaminyltransferases, and β-(1,4)-galactosyltransferases, enzymes known for their role of adding fucose and galactose sugars in protein-glycosylation and glycan chain construction (Almeida, et al. 1997; de Vries, et al. 2001; Ma, et al. 2006; Li, et al. 2018; Rini, et al. 2022). Host glycans may be used by spores, from parasites such as the bacterial pathogen *Helicobacter pylori*, for attachment. FuT, the most prevalent kind of glycosyltransferases in the PRC and in *Daphnia* genomes, catalyzes the final step in glycan chain biosynthesis by transferring fucose sugars to peripheral positions of the glycan, potentially influencing pathogen recognition and adhesion. Interestingly, α-(1,4)-N-acetylglucosaminyltransferase, which is essential for the biosynthesis of O-linked oligosaccharides (O-glycans) in gastric mucosa, plays an essential role in inhibiting *H. pylori* in mice (Kawakubo, et al. 2004). Mutant mice who lack this gene exhibit remodeling of the O-glycans that leads to elevated infection rates by *H. pylori* (Karasawa, et al. 2012). Overall, the identification of FuT islands in the PRC highlights a promising avenue for understanding the molecular mechanisms that underlie parasite resistance evolution in *D. magna*. However, it is crucial first to validate the functional role of FuT and GalT genes in parasite attachment.

Our study suggests that the PRC resides within an exceptionally old HDR. This finding potentially positions the PRC as one of the most ancient examples of polymorphisms identified so far in animals outside the jawed vertebrates. While ancient polymorphisms have previously been documented (reviewed in (Fortier and Pritchard 2022), these primarily involve the Major Histocompatibility Complex (MHC) super-locus of jawed vertebrates, with MHC-linked polymorphism in teleost fish being around 300 Mya old (Tsukamoto, et al. 2012; Grimholt, et al. 2015). An old polymorphic region was also described in basidiomycete fungi, estimated to be about 370 Mya old (Devier, et al. 2009). Because trans-species polymorphism are rarely obtained and less studied in invertebrates (Croze, et al. 2017), our work presents an important milestone for exploring host-parasite coevolution within the context of ancient, intra- and inter-species hyperdivergent regions, extending the focus beyond vertebrates and plants and beyond classical trans-species polymorphism studies.

## Conclusions

This study unveils the extraordinary polymorphism and antiquity of the *Daphnia magna* Pasteuria Resistance Complex (PRC). The exceptional intra- and inter-species variation observed within the PRC’s hyper-divergent region suggests a crucial role for balancing selection in maintaining this diversity. The presence of multiple “FuT islands” encoding various glycosyltransferases further strengthens the link between this region and potential resistance mechanisms. The persistence of highly divergent haplotypes within the PRC underscores the effectiveness of balancing selection in maintaining functional variation within populations. It suggests a potentially widespread phenomenon where hyper-divergent regions act as reservoirs of genetic diversity, continually evolving under host-parasite co-evolution. This study paves the way for future investigations into the relationship between hyper-divergent regions and balancing selection across diverse taxa. Exploring this connection could lead to a more comprehensive understanding of how balancing selection shapes the evolution and maintenance of crucial functional units within genomes, ultimately contributing to long-term adaptation and resilience in host-pathogen interactions.

## Methods

### DNA extraction and genome sequencing

To isolate DNA for whole-genome sequencing, 14 different non-*D. magna* clones from 13 different Daphniidae species (Supplementary Table 1) were cultured from juveniles. For each clone/species, 280 individuals were raised in 12 separate jars containing artificial Daphnia medium (ADaM) at 20 °C with a 16:8 light/dark cycle. Every two days, they were fed green algae (*Tetradesmus* sp.), and jars were checked every 3-4 days to remove offspring. Medium was changed once a week.

After two weeks, 240 adult females from each culture were treated with a mixture of Streptomycin, Tetracycline, and Ampicillin antibiotics (each at 50 mg/L) for three days to eliminate gut microbes (Dukic, et al. 2019). Following this antibiotic treatment, five animals from each culture were transferred into a washing jar containing a sterile 3:4 deionized water:ADaM solution, totaling 48 washing jars for each culture. Animals from all the washing jars were collected on wet net filters (25-30 per filter), centrifuged at 4 °C and 4250 rpm, and then transferred to cryotubes, frozen in liquid nitrogen for 10 seconds, and stored at -80 °C.

DNA extraction was performed using a modified protocol based on the Qiagen Genomic DNA buffer set with Genomic-tip 100/G (Blood & Cell Culture DNA Kit, cat. no. 13343). Wide bore pipette tips were used throughout to minimize DNA fragmentation, and pipetting was avoided whenever possible, opting for careful pouring instead.

Extracted DNA was then quantified using a Qubit fluorometer High sensitivity assay for double stranded DNA. Purity was assessed using a Nanodrop to measure the 260/280 ratio (ideally greater than 1.8) and the 260/230 ratio (ideally between 2.0 and 2.2), to identify potential contamination from reagents or other contamination. Finally, fragment analysis and a second quality control step were performed to ensure the DNA was suitable for PacBio HiFi sequencing.

We employed two sequencing and respective assembly approaches: Continuous Long Reads (CLR) using the FALCON pipeline (Chin, et al. 2016) with a parameter set optimized for Daphnia genome assembly, and PacBio HiFi Sequencing. Due to the significantly lower error rate of HiFi data compared to CLR, we first employed the pbccs tool (https://github.com/PacificBiosciences/ccs) to convert error-prone subreads into high-fidelity circular consensus reads (CCSs). Subsequently, individual HiFi assemblies were generated using the hifiasm v.0.15.1-r334 assembler (Cheng, et al. 2021).

To account for potential variation in genetic diversity between *D. magna* haplotypes, we conducted two separate assemblies for HiFi data. The first assembly included the “-l0” flag, which disables automatic purging of duplications. The second assembly excluded this flag, allowing the assembler to remove putative duplications or heterozygous regions. Following assembly, AWK (Aho et al., 1988) was used (as recommended on https://github.com/chhylp123/hifiasm) to convert the resulting assembly graphs into a set of contigs in a multi-fasta format. Finally, homozygous and heterozygous assemblies were evaluated for contiguity, completeness of biological information, and the best assembly was chosen for further analysis.

In addition, 14 high-quality *D.magna* published assemblies under the same project (Supplementary Table 3) were used in this study.

### Mapping the PRC in *D. magna*

Using the PRC mapped by Bento et al. (2017) and Fredericksen et al. (2023) as a reference, we obtained the PRC from 13 additional *D. magna* clones using BLASTN v2.12.0 (Altschul, et al. 1990) with default parameters.

Using PanGraph (Noll, et al. 2023), we constructed a pan-genome graph of the PRC for 14 *D. magna* clones. Due to the highly non-homologous nature of the locus, a limited set of core pancontigs was retrieved using MMseqs2 (Steinegger and Soding 2017) as the alignment kernel coupled with null energy parameters (pangraph v0.7.3 with parameters -k mmseqs -a 0 -b 0).

In order to identify genes in those pancontigs, we used BLASTX (Camacho, et al. 2009) v 2.12.0 against the uniProt database (UniProt 2023) release-2023_05.

### Mapping the PRC in other species

To obtain the PRC in additional species, we first performed a BLASTX v2.12.0 search against the predicted PRC genes from *D. magna’s* conserved pancontigs. This analysis identified the PRC region in an additional 20 non-*D. magna* assemblies based on significant sequence similarity to known *D. magna* PRC genes.

For each identified putative PRC region, we then employed BLASTN v2.12.0 to extract the corresponding sequence from the respective non-*D. magna* assembly using the established *D. magn*a PRC sequence as a reference. If the exact boundaries of the putative PRC region were ambiguous, we iteratively extended the search by adding flanking sequences from the *D. magna* PRC region until either a significant alignment was achieved or the end of the contig in *D. magna* assembly was reached, whichever occurred first. This iterative approach enabled us to capture incomplete or divergent PRC regions in other species.

### Identifying FuT islands in the PRC region and its flanking regions

To identify all FuT and GalT copies in and around the PRC, we first used BLASTX v2.12.0 for the whole contig/chromosome containing the PRC against the uniProt database; then we used BLASTP 2.12.0+ against all the FuT and GalT genes identified from the previous BLASTX run. Since we were only interested in potentially functioning genes, we filtered out all sequences with no open reading frame using ORF Finder v0.4.3 with parameters -ml 300 - s 1 -n True, or with parameters, -ml 150 -s 1 -n True for FuT and GalT, respectively (Wheeler, et al. 2003).

### Reconstructing the species tree and the gene tree

To reconstruct the species tree, we found all arthropod BUSCO genes (v5.4.3) (Manni, et al. 2021) with parameters -m genome -l arthropoda_odb10. We then concatenated and aligned the sequences of both nucleotides and amino acids using MAFFT v7.520 (Nakamura, et al. 2018) with parameters --threadit 0 --reorder --anysymbol --leavegappyregion --maxiterate 20 --retree 1 –genafpair and trimmed the alignments using trimAl v1.4.rev15 (Capella-Gutierrez, et al. 2009) with parameters -nexus -gappyout -keepheader. Using the non-reversible model in IQTREE2 v2.0.6 (Minh, et al. 2020), we inferred the rooted species tree with 1000 ultrafast bootstraps (Minh, et al. 2013; Hoang, et al. 2018) and a separate substitution model for each gene (Chernomor, et al. 2016).

For the gene tree estimation, we used two different methods: Maximum-Likelihood (ML) inference (Felsenstein 1981) and coalescent-based analysis (Kingman 2000). For the ML method, we first concatenated the genes found on the pancontigs and then aligned and trimmed them as described for the species tree. We then used IQTREE2 with ModelFinder (Kalyaanamoorthy, et al. 2017) to find the best-fit substitution model for each gene in the concatenated gene alignment and reconstructed the unrooted gene tree with 1000 ultrafast bootstraps. The gene tree was then rooted based on the species tree.

For the coalescent method, we aligned and trimmed each individual gene separately and then inferred the unrooted ML tree for each gene as described for the concatenated gene alignment. We then applied a coalescent-based method as used in ASTRAL-III v5.7.1 (Zhang, et al. 2018) with the eight gene trees to reconstruct the coalescent-based gene tree and rooted it based on the species tree.

### Reconstructing the fucosyltransferase tree

Since multiple copies of the fucosyltransferase (FuT) gene were identified within the PRC and are candidates for the *Pasteuria* resistance polymorphism observed in at least three different loci in the PRC (Bento, et al. 2017; Fredericksen, et al. 2023) (& Eglantine’s paper), we investigated the history of losses and duplications by reconciling the FuT tree with the species tree. To do so, we first aligned all the FuT copies from all taxa in which we found PRCs using MAFFT and then inferred the ML phylogeny with IQTREE2 using alpha1,3-fucosyltransferase C [Drosophila melanogaster] gene as an outgroup. Using Treerecs v1.2 (Comte, et al. 2020), we then reconciliated the FuT gene tree with the species tree and estimated the number of gene losses and gene duplications through the species tree.

## Supplementary Material

Supplementary material is available at Molecular Biology and Evolution online.

## Acknowledgements

We thank J. Hottinger, U. Stiefel and M. Krebs for help in the laboratory and the Lausanne Genomics Technology Facility for sequencing of the samples. We appreciate insightful discussions with Louis Du Pasquier during the preparation of this manuscript. We thank members of the Ebert group for feedback on the study and the manuscript. We thank S. Zweizig for language editing. This work was supported by the Swiss National Science Foundation grant numbers 310030_188887 and 310030_219529 to D.E.

## Author Contributions

All authors conceived the study. D.E. organized and collected the samples. F.S. and P.D.F conducted the sequencing work. P.D.F produced the genome assemblies. S.N. conducted the bioinformatic analysis. S.N. wrote the manuscript, which was read, edited and approved by all authors.

## Data availability

The genomic data generated in this study have been deposited in the NCBI database under accession code BioProjectID PRJEB27855. The data generated in this study are provided in the Supplementary Information/Source Data file.

Scripts required for replicating our results are available at GitHub under the repository https://github.com/suhanaser/PRC.

