## Supplementary for "The Evolution of Extreme Genetic Variability in a Parasite-Resistance Complex in a Planktonic Crustacean"

### Supplementary Material

#### Figures

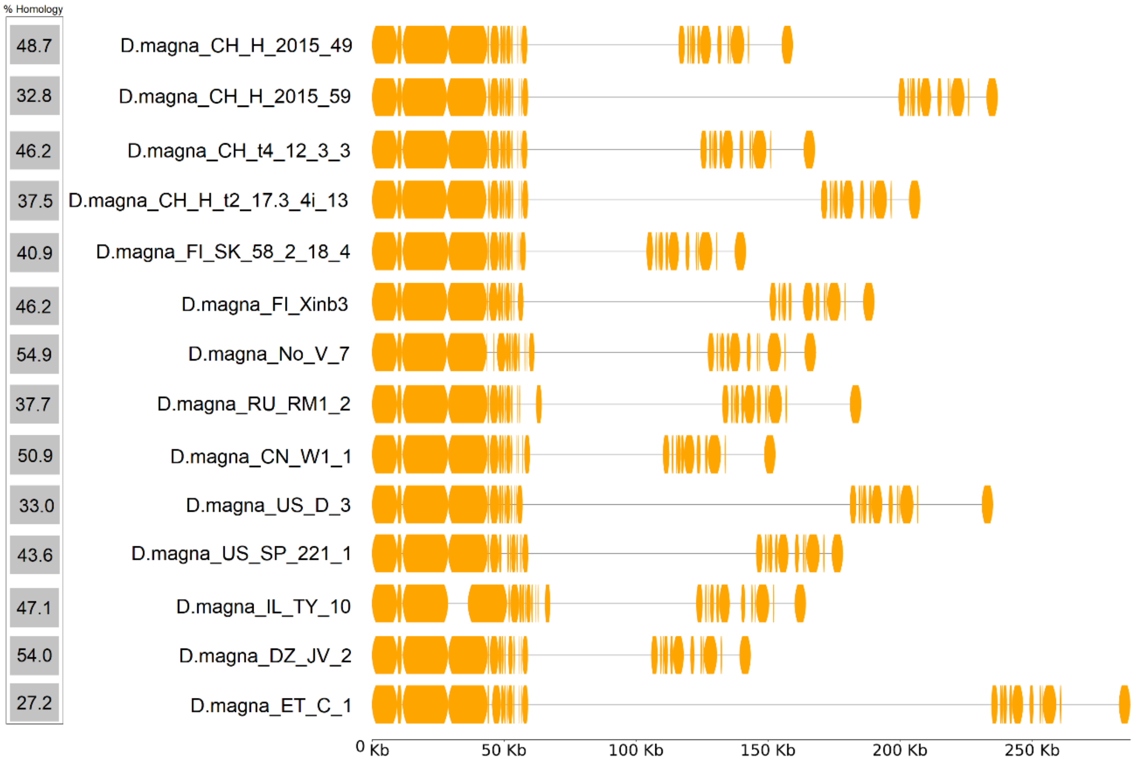


**Fig. 1.** Alignment of 14 clones of D.magna. in Orange: homologous regions.

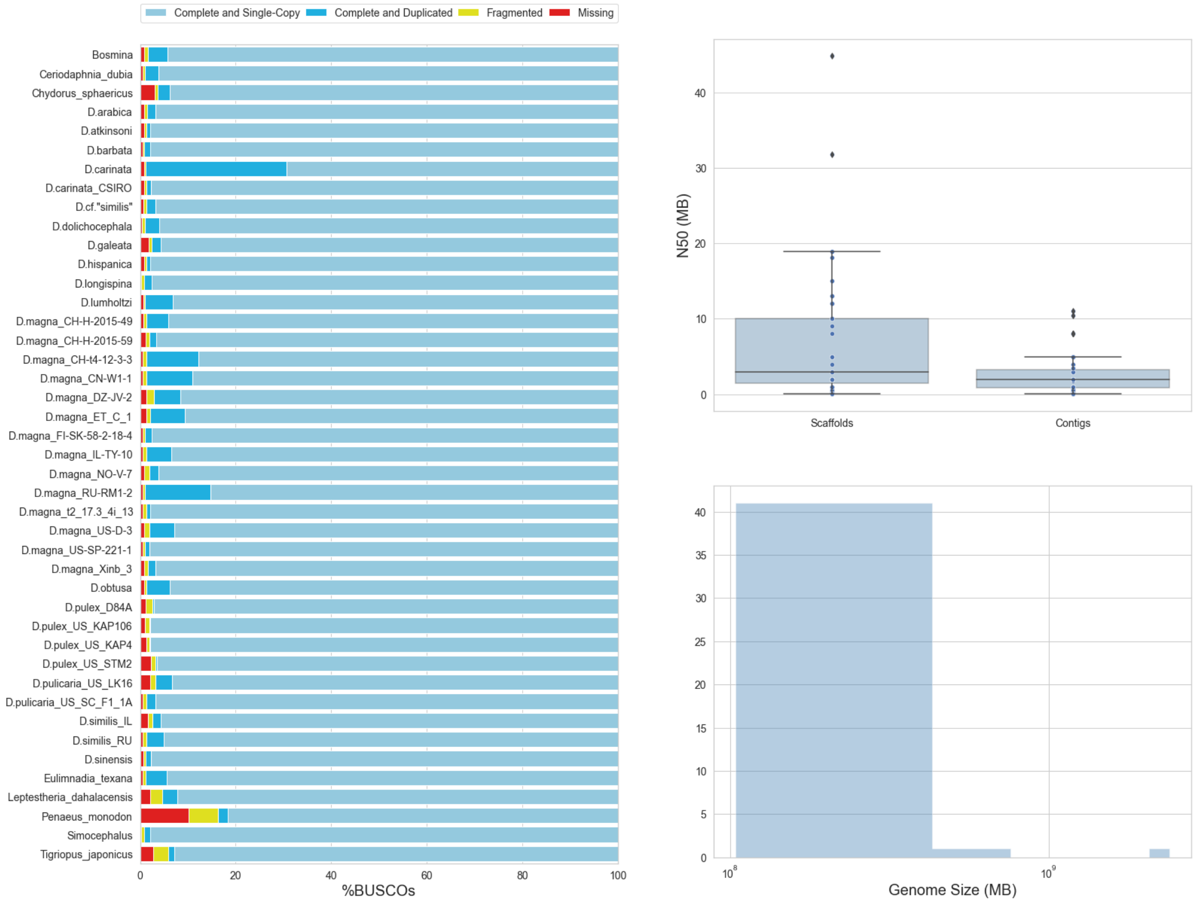


**Fig. 2.** The 43 assemblies that were used in this study. A) the Busco completeness score for each assembly. B) the distribution of N50 scaffolds and N50 contigs in MB. C) the distribution of genome size in MB.

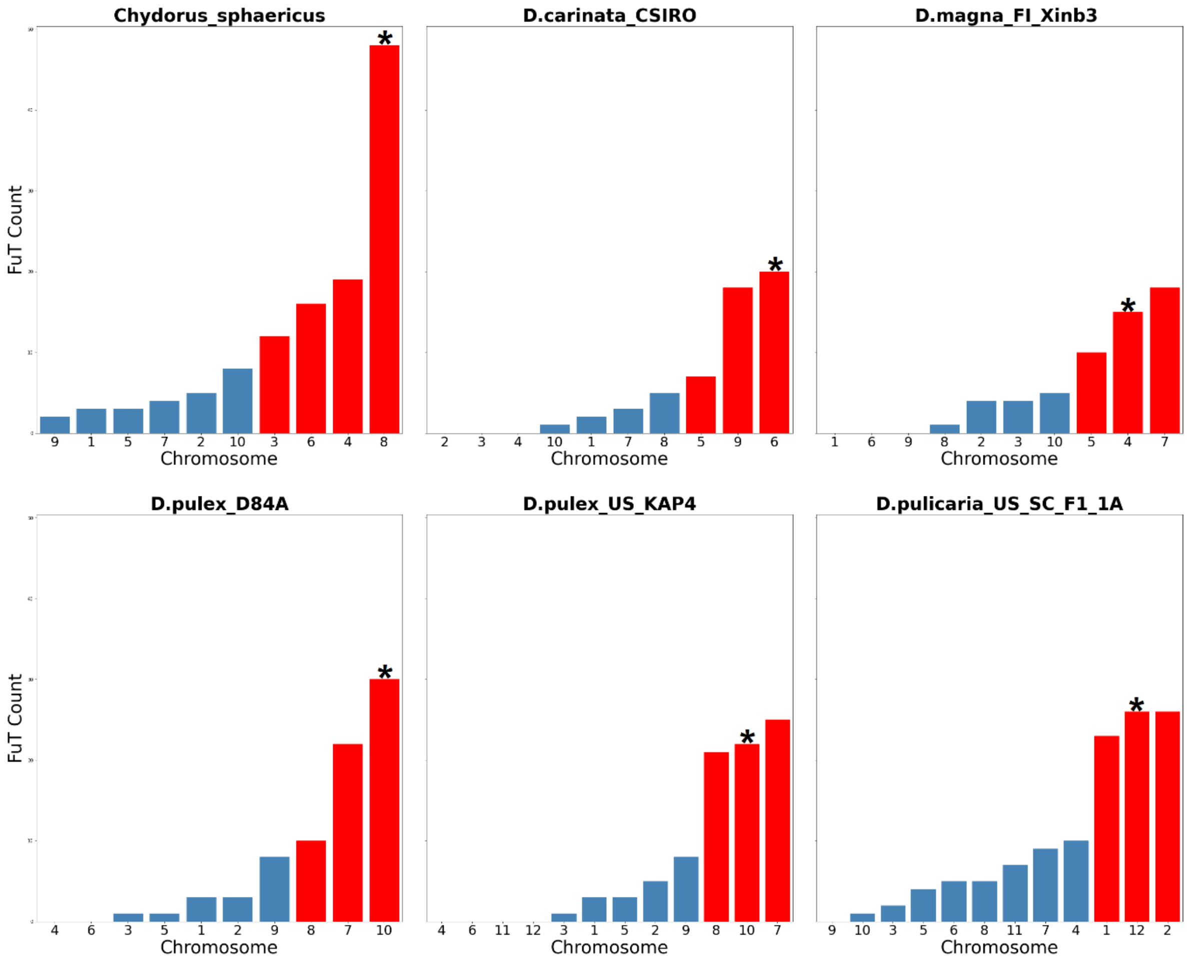


**Fig. 3.** Distribution of FuT copies across the different chromosomes in six different species. Red bars are chromosomes that together have more than 75% of the FuT copies in the genome. Asterisk is for the chromosome that contains the PRC.

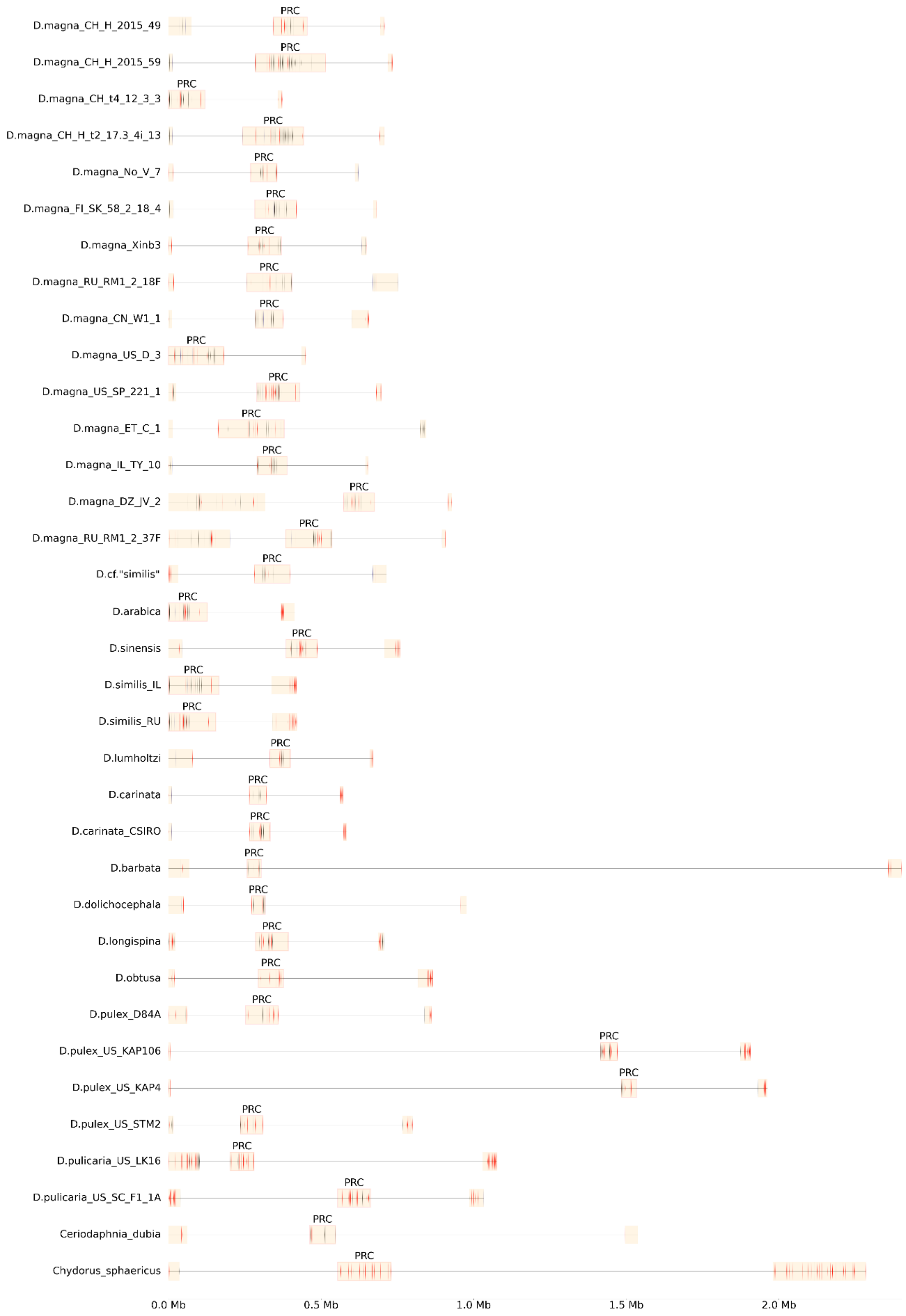


**Fig. 4.** PRC alignment of all species that have the complex and the two FucT islands surrounding the PRC.

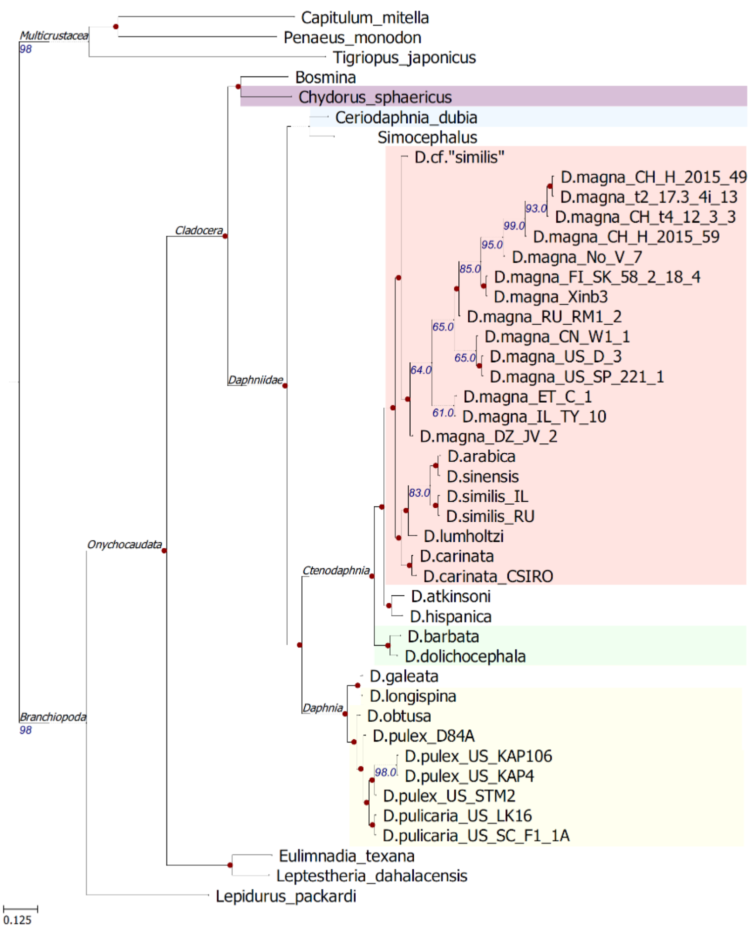


**(A)**

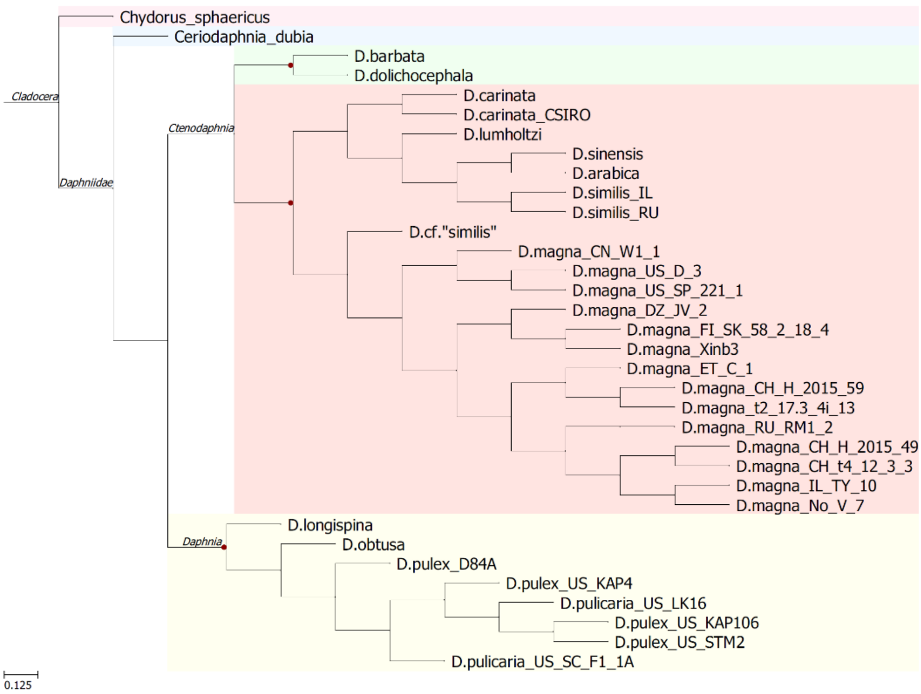


**(B)**

**Fig. 5**. A) ML phylogeny from arthropods BUSCO genes for all the assemblies used in this study B) coalescent-based phylogeny with the eight conserved PRC genes from 34 taxa

F0 clade

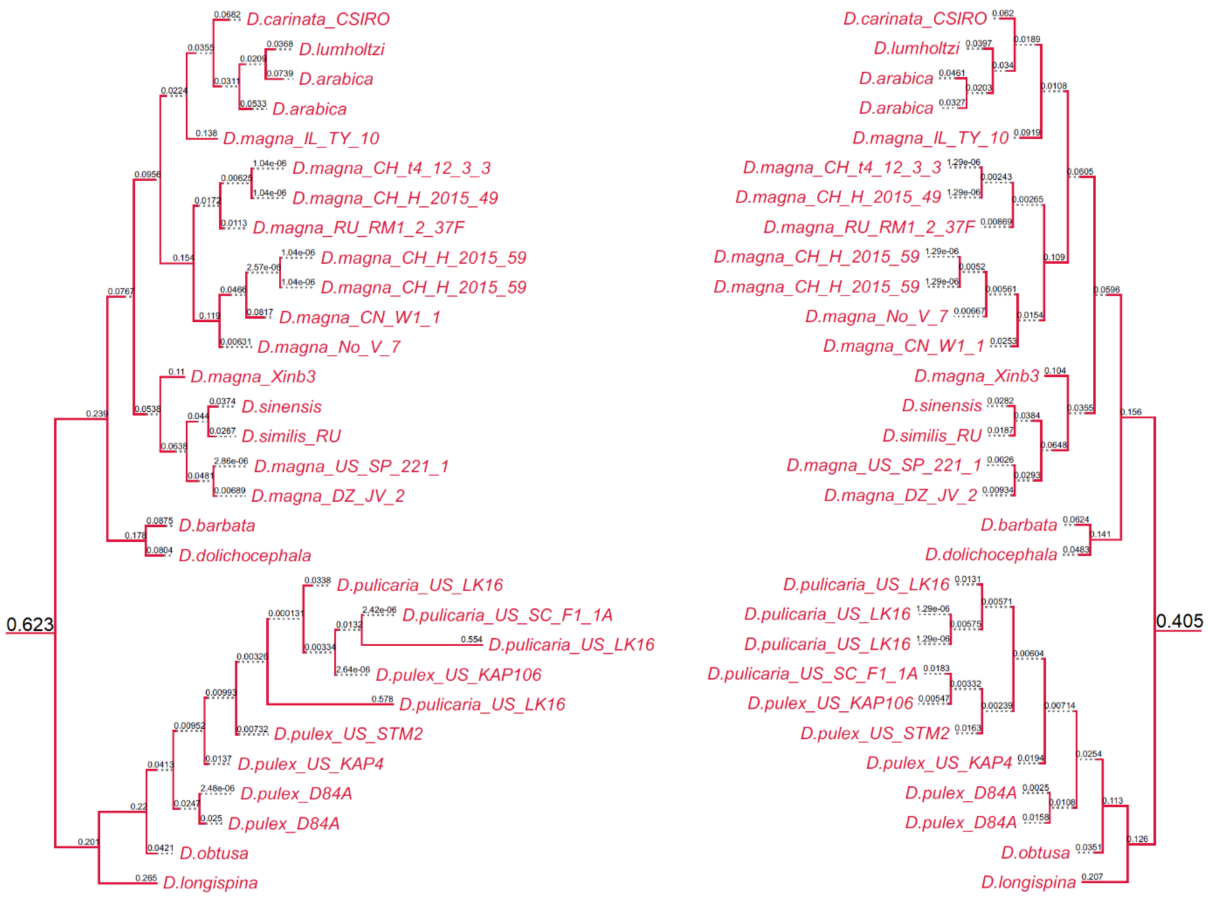
D.carinata_CSIRO
D.lumholtzi
D.arabica
D.arabica
D.magna_IL_TY_10
D.magna_CH_t4_12_3_3
D.magna_CH_H_2015_49
D.magna_RU_RM1_2_37F
D.magna_CH_H_2015_59
D.magna_CH_H_2015_59
D.magna_CN_W1_1
D.magna_No_V_7
D.magna_Xinb3
D.sinensis
D.similis_RU
D.magna_US_SP_221_1
D.magna_DZ_JV_2
D.barbata
D.dolichocephala
D.pulicaria_US_LK16
D.pulicaria_US_SC_F1_1A
D.pulicaria_US_LK16
D.pulex_US_KAP106
D.pulicaria_US_LK16
D.pulex_US_STM2
D.pulex_US_KAP4
D.pulex_D84A
D.pulex_D84A
D.obtusa
D.longispina
0.623
0.405

F1 clade

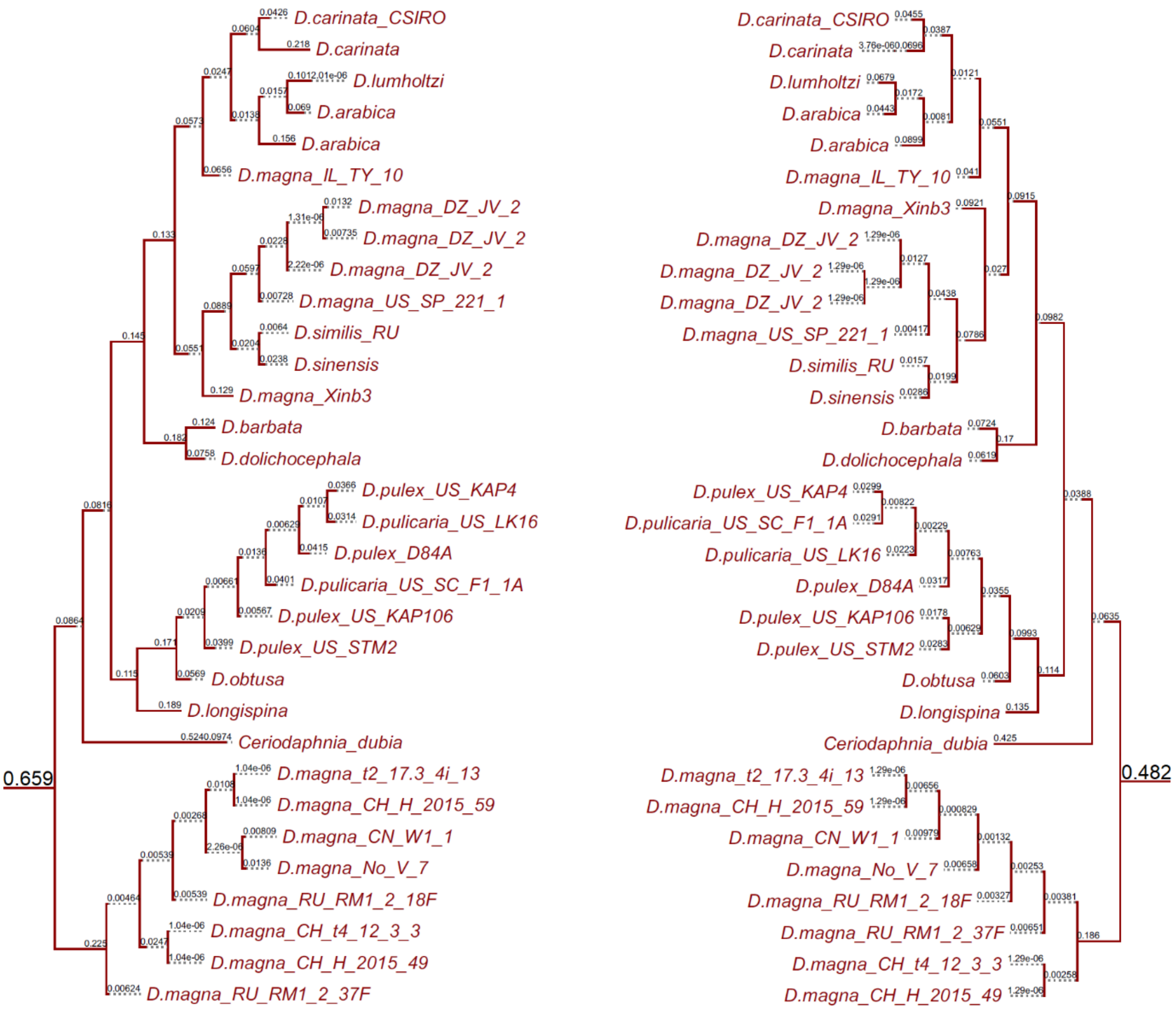
D.carinata_CSIRO
D.carinata
D.lumholtzi
D.arabica
D.arabica
D.magna_IL_TY_10
D.magna_DZ_JV_2
D.magna_DZ_JV_2
D.magna_DZ_JV_2
D.magna_US_SP_221_1
D.similis_RU
D.sinensis
D.magna_Xinb3
D.barbata
D.dolichocephala
D.pulex_US_KAP4
D.pulicaria_US_LK16
D.pulex_D84A
D.pulicaria_US_SC_F1_1A
D.pulex_US_KAP106
D.pulex_US_STM2
D.obtusa
D.longispina
Ceriodaphnia_dubia
0.659
D.magna_t2_17.3_4i_13
D.magna_CH_H_2015_59
D.magna_CN_W1_1
D.magna_No_V_7
D.magna_RU_RM1_2_18F
D.magna_CH_t4_12_3_3
D.magna_CH_H_2015_49
D.magna_RU_RM1_2_37F
D.carinata_CSIRO
D.carinata
D.lumholtzi
D.arabica
D.arabica
D.magna_IL_TY_10
D.magna_Xinb3
D.magna_DZ_JV_2
D.magna_DZ_JV_2
D.magna_DZ_JV_2
D.magna_US_SP_221_1
D.similis_RU
D.sinensis
D.barbata
D.dolichocephala
D.pulex_US_KAP4
D.pulicaria_US_SC_F1_1A
D.pulicaria_US_LK16
D.pulex_D84A
D.pulex_US_KAP106
D.pulex_US_STM2
D.obtusa
D.longispina
Ceriodaphnia_dubia
0.482
D.magna_t2_17.3_4i_13
D.magna_CH_H_2015_59
D.magna_CN_W1_1
D.magna_No_V_7
D.magna_RU_RM1_2_18F
D.magna_RU_RM1_2_37F
D.magna_CH_t4_12_3_3
D.magna_CH_H_2015_49

F2 clade

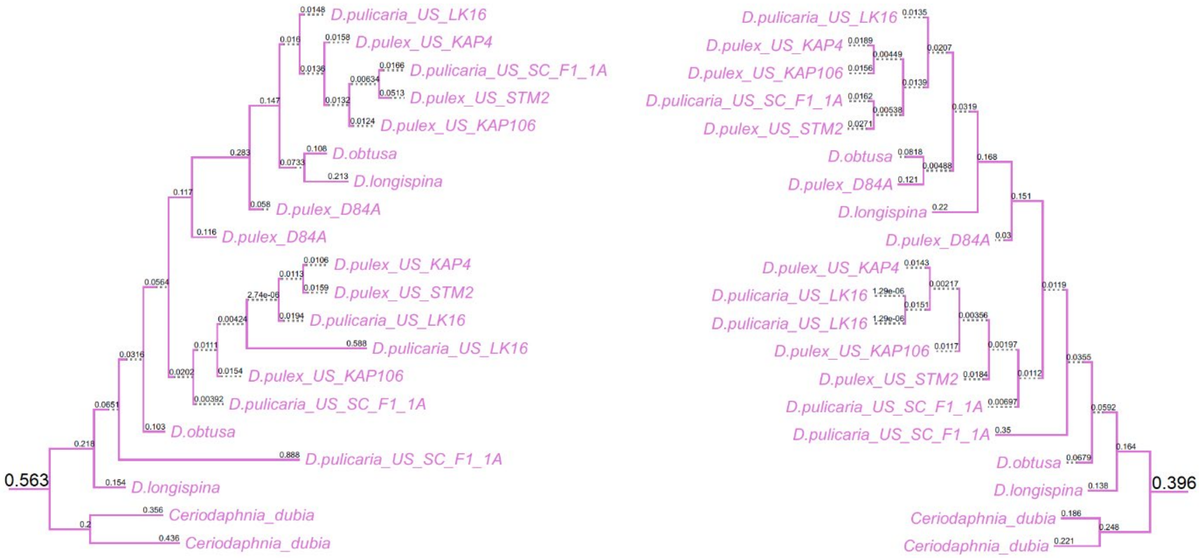
D.pulicaria_US_LK16
D.pulex_US_KAP4
D.pulicaria_US_SC_F1_1A
D.pulex_US_STM2
D.pulex_US_KAP106
D.obtusa
D.longispina
D.pulex_D84A
D.pulex_D84A
D.pulex_US_KAP4
D.pulex_US_STM2
D.pulicaria_US_LK16
D.pulicaria_US_LK16
D.pulex_US_KAP106
D.pulicaria_US_SC_F1_1A
D.obtusa
D.pulicaria_US_SC_F1_1A
D.longispina
Ceriodaphnia_dubia
Ceriodaphnia_dubia
0.563
D.pulicaria_US_LK16
D.pulex_US_KAP4
D.pulex_US_KAP106
D.pulicaria_US_SC_F1_1A
D.pulex_US_STM2
D.obtusa
D.pulex_D84A
D.longispina
D.pulex_D84A
D.pulex_US_KAP4
D.pulicaria_US_LK16
D.pulicaria_US_LK16
D.pulex_US_KAP106
D.pulex_US_STM2
D.pulicaria_US_SC_F1_1A
D.pulicaria_US_SC_F1_1A
D.obtusa
D.longispina
Ceriodaphnia_dubia
Ceriodaphnia_dubia
0.396

F3 clade

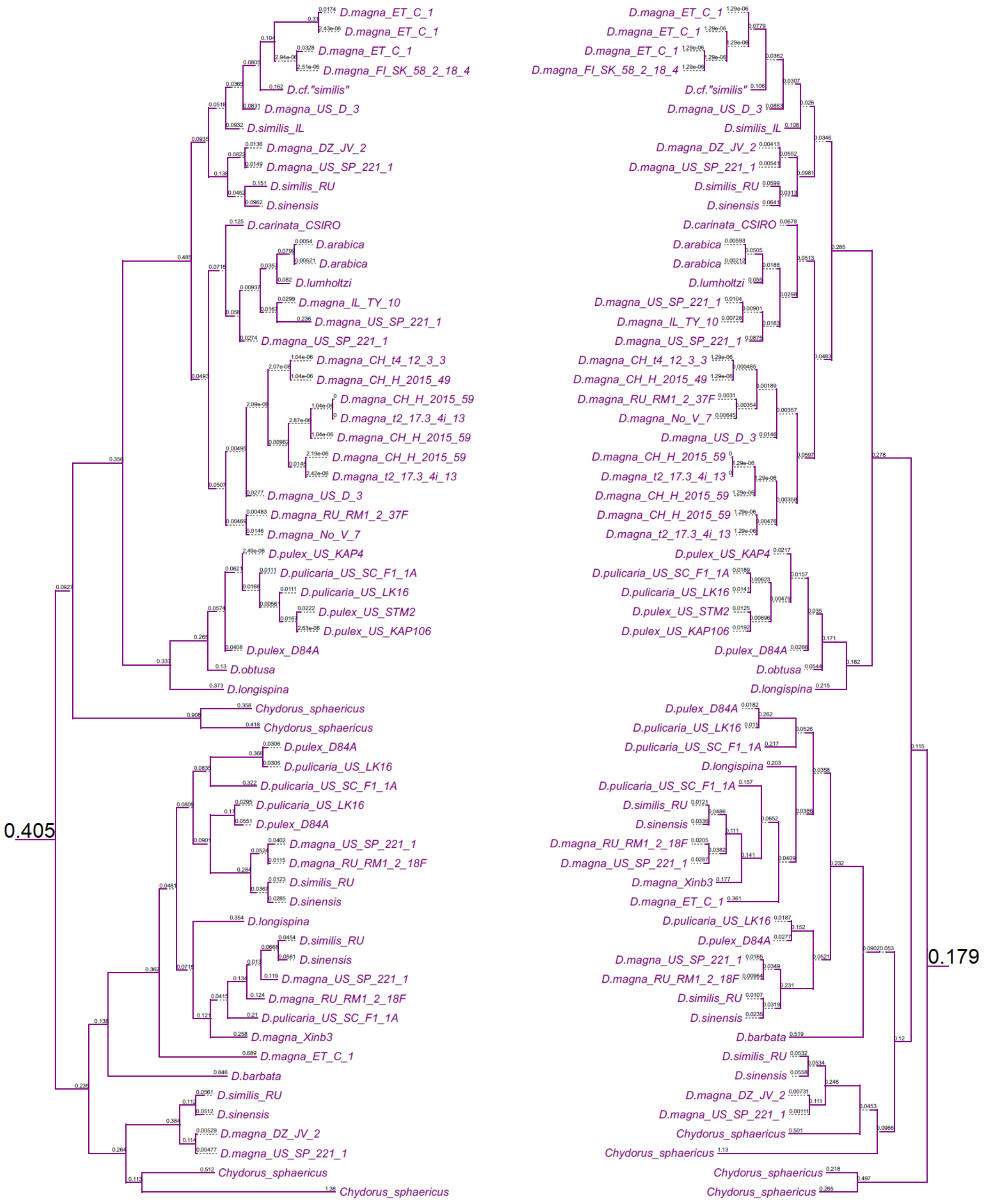
0.405
0.179
D.magna_ET_C_1
D.magna_ET_C_1
D.magna_ET_C_1
D.magna_FI_SK_58_2_18_4
D.cf."similis"
D.magna_US_D_3
D.similis_IL
D.magna_DZ_JV_2
D.magna_US_SP_221_1
D.similis_RU
D.sinensis
D.carinata_CSIRO
D.arabica
D.arabica
D.lumholtzi
D.magna_IL_TY_10
D.magna_US_SP_221_1
D.magna_US_SP_221_1
D.magna_CH_t4_12_3_3
D.magna_CH_H_2015_49
D.magna_CH_H_2015_59
D.magna_t2_17.3_4i_13
D.magna_CH_H_2015_59
D.magna_CH_H_2015_59
D.magna_t2_17.3_4i_13
D.magna_US_D_3
D.magna_RU_RM1_2_37F
D.magna_No_V_7
D.pulex_US_KAP4
D.pulicaria_US_SC_F1_1A
D.pulicaria_US_LK16
D.pulex_US_STM2
D.pulex_US_KAP106
D.pulex_D84A
D.obtusa
D.longispina
Chydorus_sphaericus
Chydorus_sphaericus
D.pulex_D84A
D.pulicaria_US_LK16
D.pulicaria_US_SC_F1_1A
D.pulicaria_US_LK16
D.pulex_D84A
D.magna_US_SP_221_1
D.magna_RU_RM1_2_18F
D.similis_RU
D.sinensis
D.longispina
D.similis_RU
D.sinensis
D.magna_US_SP_221_1
D.magna_RU_RM1_2_18F
D.pulicaria_US_SC_F1_1A
D.magna_Xinb3
D.magna_ET_C_1
D.barbata
D.similis_RU
D.sinensis
D.magna_DZ_JV_2
D.magna_US_SP_221_1
Chydorus_sphaericus
Chydorus_sphaericus

F4 clade

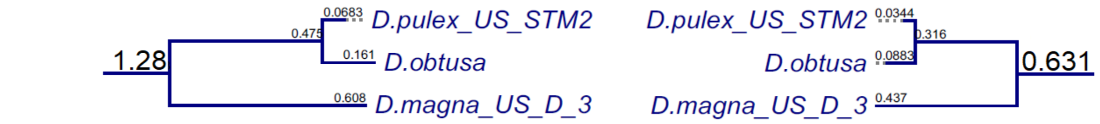


F5 clade

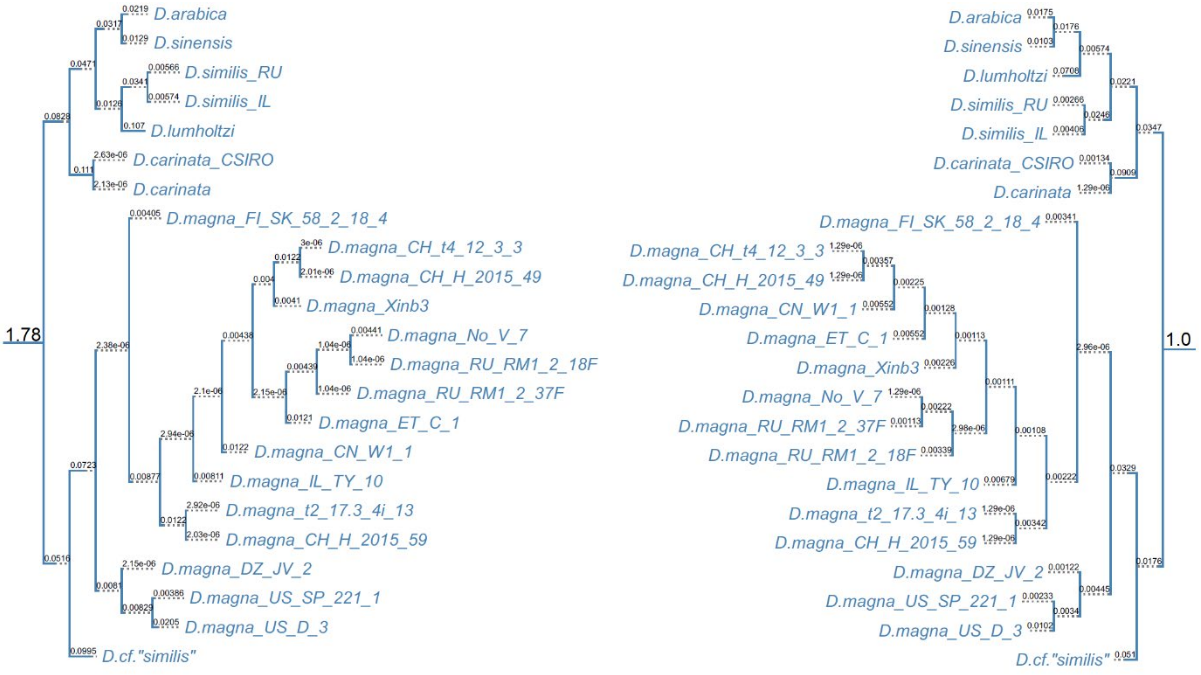

F6 clade

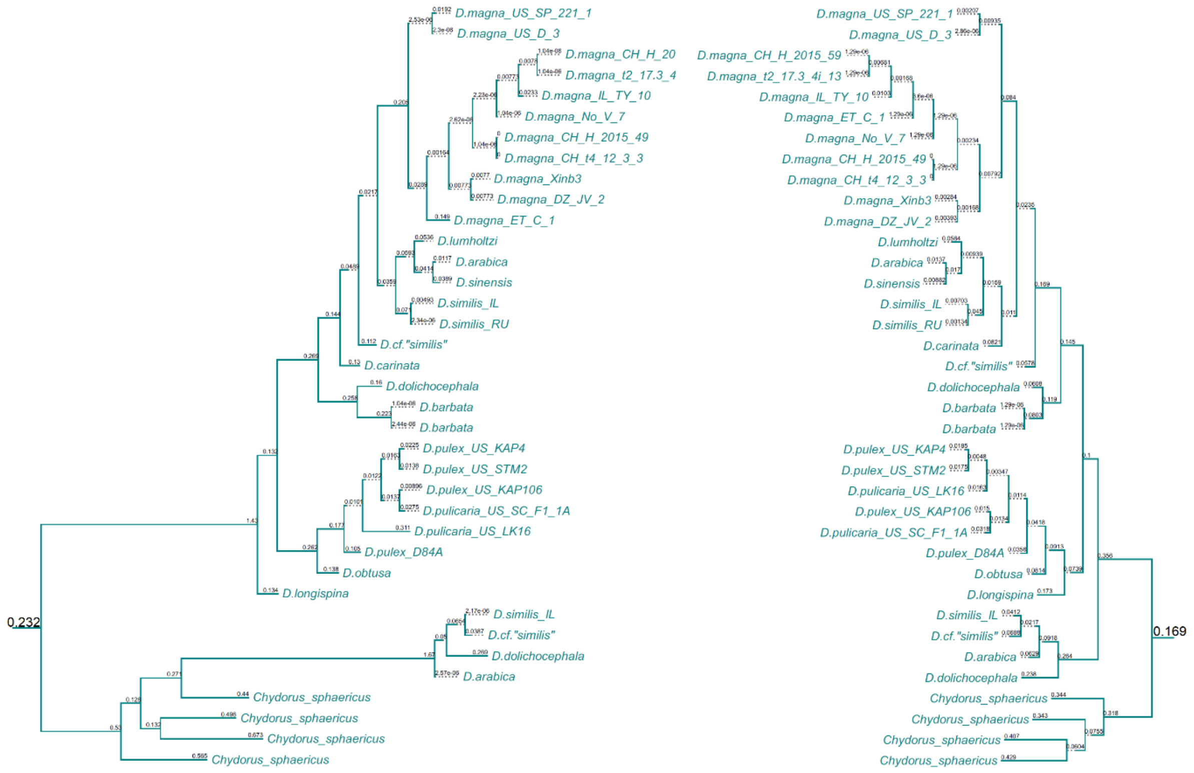


F7 clade

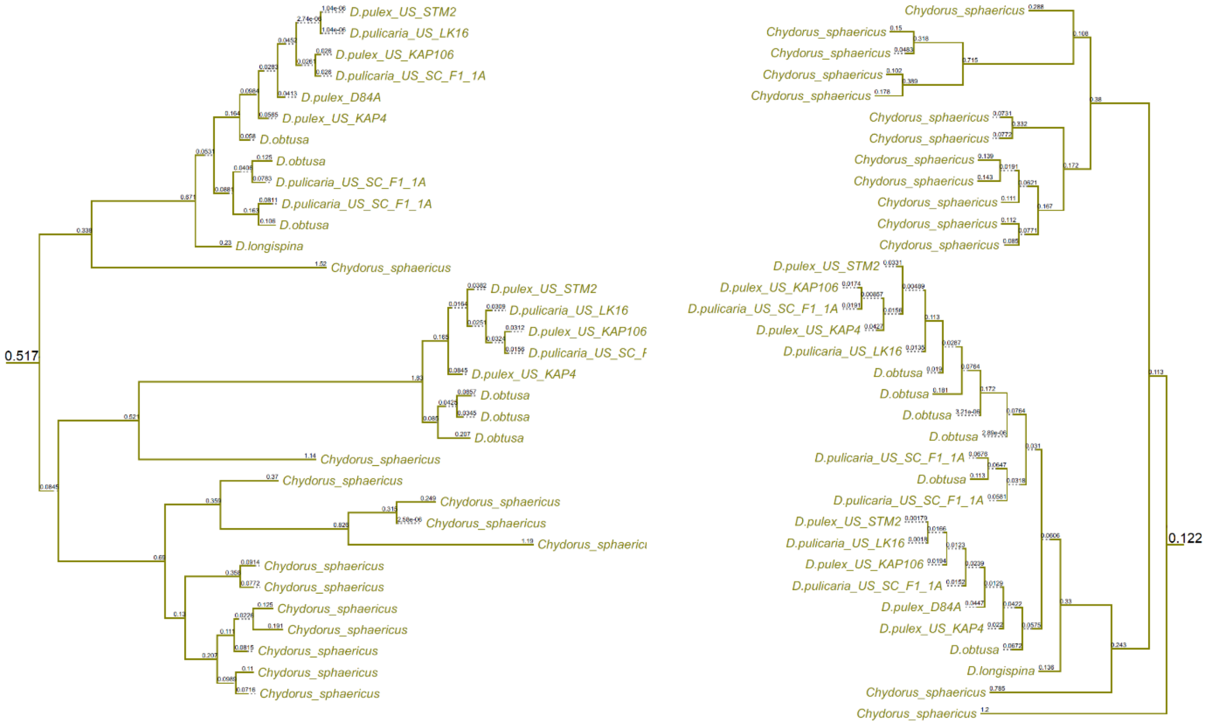


**Fig. 6.** The ML gene trees of the 8 FuT clades. Values on branches are the branch lengths. Right: trees from amino-acid alignments. Left: trees from nucleotide alignments.

### Tables

**Table 1|** The proportion of homologous sequences in the PRC between pairs of *D.magna*.

The value in cell *i,j* is the length of the homologous sequence between *i* and *j* divided by the length of *i* multiplied by 100.

Since the proportion of homology is calculated based on the length of the PRC region for the clone in the row, the upper are the lower diagonals are different.

| | CH_H_2015_49 | CH_H_2015_59 | CH_t4_12_3_3 | CN_W1_1 | ET_C_1 | DZ_JV_2 | FI_SK_58_2_18_4 | IL_TY_10 | No_V_7 | RU_RM1_2 | US_D_3 | US_SP_221_1 | Xinb3 | t2_17.3_4i_13 | **mean** |
|---|---|---|---|---|---|---|---|---|---|---|---|---|---|---|---|
| CH_H_2015_49 | | 64 | 93 | 70 | 61 | 59 | 58 | 68 | 78 | 71 | 69 | 63 | 61 | 65 | **68** |
| CH_H_2015_59 | 43 | | 43 | 46 | 42 | 39 | 39 | 41 | 44 | 39 | 39 | 39 | 41 | 55 | **42** |
| CH_t4_12_3_3 | 88 | 61 | | 66 | 58 | 56 | 55 | 64 | 74 | 68 | 64 | 58 | 58 | 61 | **64** |
| CN_W1_1 | 73 | 71 | 73 | | 60 | 58 | 62 | 61 | 76 | 71 | 68 | 60 | 60 | 71 | **66** |
| DZ_JV_2 | 68 | 69 | 68 | 63 | | 65 | 66 | 69 | 69 | 62 | 65 | 87 | 76 | 69 | **69** |
| ET_C_1 | 33 | 32 | 33 | 31 | 32 | | 39 | 33 | 34 | 32 | 33 | 33 | 32 | 33 | **33** |
| FI_SK_58_2_18_4 | 49 | 49 | 49 | 50 | 50 | 58 | | 52 | 52 | 53 | 46 | 47 | 54 | 52 | **51** |
| IL_TY_10 | 66 | 59 | 65 | 56 | 60 | 58 | 60 | | 63 | 56 | 58 | 62 | 60 | 59 | **60** |
| No_V_7 | 88 | 74 | 88 | 82 | 70 | 70 | 69 | 73 | | 76 | 73 | 70 | 72 | 74 | **75** |
| RU_RM1_2 | 49 | 40 | 49 | 47 | 38 | 40 | 43 | 39 | 46 | | 44 | 39 | 39 | 40 | **43** |
| US_D_3 | 47 | 40 | 45 | 44 | 40 | 40 | 37 | 40 | 44 | 44 | | 46 | 39 | 40 | **42** |
| US_SP_221_1 | 56 | 52 | 55 | 51 | 70 | 53 | 50 | 57 | 56 | 51 | 61 | | 56 | 52 | **55** |
| Xinb3 | 58 | 58 | 58 | 55 | 64 | 55 | 61 | 58 | 61 | 54 | 55 | 60 | | 58 | **58** |
| t2_17.3_4i_13 | 50 | 63 | 49 | 52 | 48 | 45 | 47 | 46 | 50 | 45 | 45 | 45 | 47 | | **49** |
| **mean** | **59** | **56** | **59** | **55** | **53** | **54** | **53** | **54** | **57** | **56** | **55** | **55** | **53** | **56** | **55** |

**Table 2|** Summary statistics of the Pasteuria ramosa complex FucT and of the to FucT islands to the left and right of it.

| **Mean** | **Left FucT island** | **PRC FucT island** | **Right FucT island** |
|---|---|---|---|
| No. of FucT + GalT copies | 9 | 20 | 12 |
| No. of FucT copies | 5 | 10 | 9 |
| distance between two adjacent copies | 5,094 bp | 5,553 bp | 4,062 bp |
| distance from PRC | 343,437 bp | | 407,374 bp |
| **Max** | | | |
| No. of FucT + GalT copies | 30 | 52 | 64 |
| No. of FucT copies | 20 | 33 | 63 |
| distance between copies | 43,549 bp | 66,672 bp | 42,765 bp |
| distance from PRC | 1,476,796 bp | | 2,053,738 bp |
| **Min** | | | |

| | | | |
|---|---|---|---|
| No. of FucT + GalT copies | 3 | 9 | 3 |
| No. of FucT copies | 1 | 3 | 1 |
| distance from PRC | 99,213 bp | | 172,348 bp |

**Median**

| | | | |
|---|---|---|---|
| No. of FucT + GalT copies | 6 | 16 | 8 |
| No. of FucT copies | 4 | 9 | 7 |
| distance between copies | 946 bp | 1,969 bp | 580 bp |
| distance from PRC | 255,862 bp | | 259,410 bp |

**Table 3|** Genome assemblies used in this study

| Taxon name | Species | BioProject |
|---|---|---|
| **Bosmina** | Bosmina (Eubosmina) coregoni | PRJEB27855 |
| **Capitulum_mitella** | Capitulum mitella | PRJNA816681 |
| **Ceriodaphnia_dubia** | Ceriodaphnia dubia | PRJNA798169 |
| **Chydorus_sphaericus** | Chydorus sphaericus | PRJNA967142 |
| **D.CA_CBC** | D.cf."similis" | PRJEB27855 |
| **D.arabica** | Daphnia arabica | PRJNA904511 |
| **D.atkinsoni** | Daphnia atkinsoni | PRJEB27855 |
| **D.barbata** | Daphnia barbata | PRJEB27855 |
| **D.carinata** | Daphnia carinata | PRJEB27855 |
| **D.carinata_CSIRO** | Daphnia carinata | PRJNA798159 |
| **D.dolichocephala** | Daphnia dolichocephala | PRJEB27855 |
| **D.galeata** | Daphnia galeata | PRJNA985241 |
| **D.hispanica** | Daphnia hispanica | PRJEB27855 |
| **D.longispina** | Daphnia longispina | PRJEB27855 |
| **D.lumholtzi** | Daphnia lumholtzi | PRJEB27855 |
| **D.magna_CH_H_2015_49** | Daphnia magna | PRJEB27855 |
| **D.magna_CH_H_2015_59** | Daphnia magna | PRJEB27855 |
| **D.magna_CH_t4_12_3_3** | Daphnia magna | PRJEB27855 |
| **D.magna_CN_W1_1** | Daphnia magna | PRJEB27855 |
| **D.magna_DZ_JV_2** | Daphnia magna | PRJEB27855 |
| **D.magna_ET_C_1** | Daphnia magna | PRJEB27855 |
| **D.magna_FI_SK_58_2_18_4** | Daphnia magna | PRJEB27855 |
| **D.magna_IL_TY_10** | Daphnia magna | PRJEB27855 |
| **D.magna_No_V_7** | Daphnia magna | PRJEB27855 |
| **D.magna_RU_RM1_2** | Daphnia magna | PRJEB27855 |
| **D.magna_US_D_3** | Daphnia magna | PRJEB27855 |
| **D.magna_US_SP_221_1** | Daphnia magna | PRJEB27855 |
| **D.magna_Xinb3** | Daphnia magna | PRJEB27855 |
| **D.magna_t2_17.3_4i_13** | Daphnia magna | PRJEB27855 |
| **D.obtusa** | Daphnia obtusa | PRJNA598691 |

| | | |
|---|---|---|
| **D.pulex_D84A** | Daphnia pulex EU | PRJNA725506 |
| **D.pulex_US_KAP106** | Daphnia pulex US | PRJNA777597 |
| **D.pulex_US_KAP4** | Daphnia pulex US | PRJNA777597 |
| **D.pulex_US_STM2** | Daphnia pulex US | PRJNA777597 |
| **D.pulicaria_US_LK16** | Daphnia pulicaria | PRJNA686130 |
| **D.pulicaria_US_SC_F1_1A** | Daphnia pulicaria | PRJNA762351 |
| **D.similis_IL** | Daphnia similis | PRJEB27855 |
| **D.similis_RU** | Daphnia similis | PRJEB27855 |
| **D.sinensis** | Daphnia sinensis | PRJEB27855 |
| **Eulimnadia_texana** | Eulimnadia texana | PRJNA352082 |
| **Lepidurus_packardi** | Lepidurus packardi | PRJNA811174 |
| **Leptestheria_dahalacensis** | Leptestheria dahalacensis | PRJNA417576 |
| **Penaeus_monodon** | Penaeus monodon | PRJNA611030 |
| **Simocephalus** | Simocephalus serrulatus | PRJEB27855 |
| **Tigriopus_japonicus** | Tigriopus japonicus | PRJNA592403 |

**Table 4|** The number of duplications and losses of each clade of FucT along the species tree

| Clade | Duplications | Losses |
|---|---|---|
| F0 | 4 | 20 |
| F1 | 8 | 39 |
| F2 | 14 | 41 |
| F3 | 10 | 81 |
| F4 | 16 | 74 |
| F5 | 0 | 12 |
| F6 | 8 | 36 |
| F7 | 17 | 70 |
| F8 | 21 | 32 |
| F9 | 12 | 56 |
| **Total** | **110** | **461** |
